# DNA binding and bridging by human CtIP in the healthy and diseased states

**DOI:** 10.1101/2023.12.14.571649

**Authors:** Shreya Lokanathan Balaji, Sara De Bragança, Francisco Balaguer-Pérez, Sarah Northall, Oliver Wilkinson, Clara Aicart-Ramos, Neeleema Seetaloo, Frank Sobott, Fernando Moreno-Herrero, Mark Simon Dillingham

## Abstract

The human DNA repair factor CtIP helps to initiate the resection of double-stranded DNA breaks for repair by homologous recombination, in part through its ability to bind and bridge DNA molecules. However, CtIP is a natively disordered protein that bears no apparent similarity to other DNA-binding proteins and so the structural basis for these activities remains unclear. In this work, we have used bulk DNA binding, single molecule tracking, and DNA bridging assays to study wild-type and variant CtIP proteins to better define the DNA binding domains and the effects of mutations associated with inherited human disease. Our work identifies a monomeric DNA-binding domain in the C-terminal region of CtIP. CtIP binds non-specifically to DNA and can diffuse over thousands of nucleotides. CtIP-mediated bridging of distant DNA segments is observed in single-molecule magnetic tweezers experiments. However, we show that binding alone is insufficient for DNA bridging, which also requires tetramerization via the N-terminal domain. Variant CtIP proteins associated with Seckel and Jawad syndromes display impaired DNA binding and bridging activities. The significance of these findings in the context of facilitating DNA break repair is discussed.

**Significance Statement:** CtIP helps to repair broken chromosomes through its ability to bind and bridge DNA molecules. We studied the structural and biochemical basis for these activities and how they are affected by hereditary CtIP mutations associated with developmental disorders. We discovered a minimal domain in the C-terminal region of CtIP which supports DNA binding as a monomer. DNA binding is non-specific and facilitates 1D diffusion, but binding alone is insufficient for intermolecular tethering of DNA molecules which requires tetramerization of CtIP via N-terminal coiled-coil domains. All disease variants tested displayed impaired DNA bridging activity. These results have important implications for understanding the role of CtIP as a hub protein for DNA break repair and its dysfunction in human disease.

## Introduction

The human DNA repair factor CtIP was originally discovered as an interaction partner for the CtBP transcriptional co-repressor but is now best-characterised for its important role in the initiation of double-stranded DNA break (DSB) repair (1, 2). CtIP dysfunction is implicated in cancer and the protein has been described as both a tumour suppressor and an oncogene in different contexts (3, 4). CtIP is mutated in Jawad and Seckel-2 syndromes which are recessively inherited diseases characterised by neurodevelopmental defects and microcephaly. These syndromes are caused by exonic frameshift and alternative splice site mutations respectively, thereby producing C-terminally truncated forms of the protein (**Figure 1A**) (5). However, early onset breast cancer and a Seckel-like disease have also been reported to be associated with a point mutation (R100W) in the N-terminal region of the protein (6, 7). Interestingly, in cell lines derived from Seckel-2 patients, full length CtIP is present in cells at normal levels but is accompanied by the variant which lacks the C-terminal domain and acts in a dominant negative manner to inhibit DSB repair (5).

**Figure 1.**
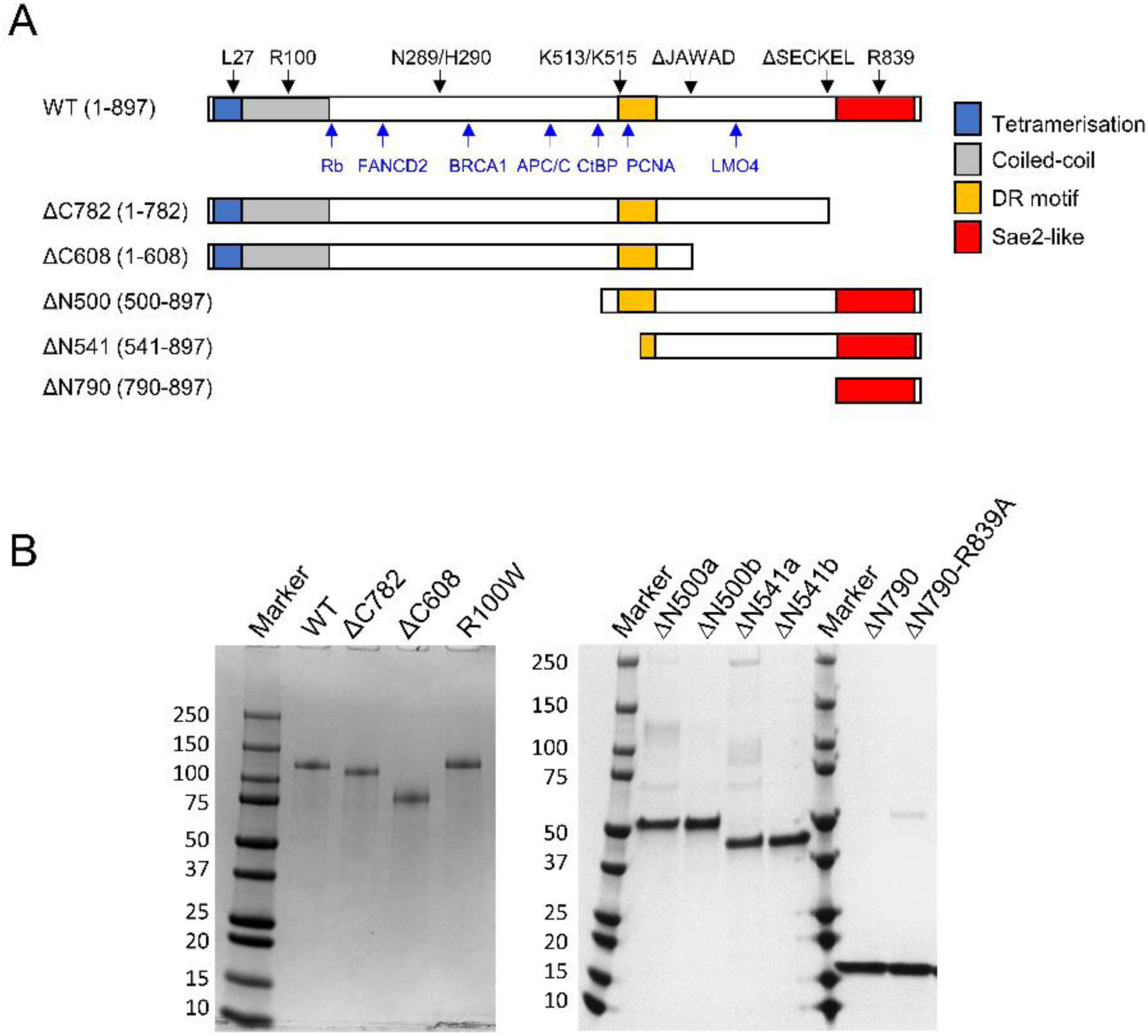
CtIP domain organisation and constructs used in this study. (A) Primary structure diagrams for wild type and CtIP deletion variants used in this study coloured according to domain organisation. The wild type sequence also indicates the positions of key amino acids and protein interaction sites above and below the sequence bar respectively. See main text for more information. (B) Purified proteins used in this study analysed by SDS-PAGE. The suffixes a and b indicate different SEC fractions of the same preparation (see text for details).

In eukaryotic cells, DSB repair proceeds through multiple degenerate pathways including homologous recombination (HR) which requires an additional DNA molecule to act as a template for faithful repair. The complex DNA transactions which promote HR require co-ordination of the two free DNA ends originating from the break site with a homology donor molecule; typically, the sister chromatid (8, 9). The process is initiated by the resection of DSBs via the collective action of DNA binding proteins, helicases and nucleases, leading to the formation of long 3ʹ-terminated ssDNA overhangs. These are substrates for RAD51-mediated homology search, strand invasion and subsequent steps of recombinational repair (8). Resection is a key control step for DSB repair, as extensive resection in the absence of a suitable donor template DNA can result in genomic instability. Consequently, DSB resection is tightly-regulated and promoted during the S/G2 phases of the cell cycle. Nevertheless, DSB resection can also play a role in initiating alternative, highly-mutagenic, DSB repair that does not use a homology donor including the single-strand annealing and micro-homology-mediated end joining pathways (10).

CtIP plays an important and multifaceted role in DSB resection but remains poorly characterised at the molecular level (for reviews see (11-13)). It is involved in HR activation in S and G2 phases of the cell cycle through CDK- and ATM-dependent phosphorylation of the S327, T847 and T859 residues (14-17), acting as a component of the BRCA1-C complex (18-22). Together, activated CtIP and MRN recognise and process “complex” DSBs in preparation for long range resection by removing tightly- or covalently-bound proteins or nucleic acid secondary structures (23-26). CtIP also promotes long range resection by recruiting and/or activating additional factors including EXO1 and DNA2 at the broken DNA (27). Importantly, CtIP is itself a DNA binding protein that can bridge DNA molecules *in trans* (28, 29), a property it shares with the MRN complex and which may be relevant to maintaining proximity between two broken DNA ends or in mediating interactions between broken DNA and the homology donor (30, 31).

The structural basis for DNA binding and bridging by CtIP is not understood. The protein forms a stable tetramer in a dimer-of-dimers arrangement with an overall architecture that resembles a dumbbell, and which can bind tightly to at least two molecules of DNA (28, 32). Each monomer comprises an N-terminal tetramerization domain (NTD) neighbouring a long coiled-coil region (32, 33), a central disordered region that is responsible for many protein-protein interactions, and a C-terminal domain (CTD) which is conserved in the yeast orthologue Sae2 (**Figure 1A**). Bioinformatics analyses of CtIP primary structure do not reveal any obvious DNA binding domains, but experimental studies have implicated three regions of the protein in the interactions with DNA. Firstly, substitution of R839 in a conserved “RHR motif” within the Sae2-like domain reduces DNA binding and bridging in full length CtIP (28, 34, 35). Secondly, a “Damage Recruitment Motif” (DR motif; residues 509-557 within the central disordered region) was identified by screening deletion mutants of CtIP fused to GFP for their translocation to laser-induced DNA damage (36). This region of the protein alone was capable of binding directly to DNA in a manner dependent on lysine residues K513 and K515. Finally, it has been suggested that residues N289 and H290 form an endonuclease active site within the central disordered region, again implicating it in DNA recognition (37). Based on the limited structural and biochemical data available, crude models have been proposed for CtIP-dependent DNA binding and bridging which envision the two DNA binding sites occupying either end of the ∼30 nm dumbbell, separated by the long coiled-coil. Together, these observations suggested to us that the N-terminal coiled-coil region of CtIP may be dispensable for DNA binding, although its role in tetramerising the protein might be pivotal in bridging DNA segments *in trans*.

In this study, we investigate the structural basis for CtIP oligomerisation, DNA binding and DNA bridging using both systematically designed N-terminal deletion constructs and disease-associated variants (**Figure 1A**). Our work demonstrates that oligomerisation of CtIP is mediated by both the NTD and central regions of the protein, and that the protein contains a monomeric DNA binding domain in the CTD. CtIP binds non-specifically to DNA and diffuses over thousands of nucleotides, facilitating the bridging between distant parts of DNA. Efficient CtIP-dependent DNA bridging requires both DNA binding and tetramerization and is impaired in all the disease variants tested. The sliding and bridging function of CtIP may underpin the co-ordination of DNA ends and donor DNA during DNA break resection and subsequent steps of HR.

## Results

### Design and purification of CtIP variants

To investigate how deletions of the CtIP C-terminal region affect activity, we purified two variant proteins, named ΔC782 and ΔC608, in which the final 115 and 289 amino-acid residues are removed from the protein respectively (**Figure 1A**). These were designed to be equivalent to the CtIP deletions that occur in Seckel-2 and Jawad syndromes. We also produced the point mutant R100W which is associated with Seckel-like disease. Finally, we made three proteins with deletions of the N-terminal region (named ΔN500, ΔN541 and ΔN790) which remove the first 499, 540 and 789 amino acids of the protein respectively. The constructs were all expressed and purified from insect cells except for ΔN790 which was well-expressed in *E. coli* (**Figure 1B**).

### The CtIP disease variants tetramerise but cannot bind to DNA efficiently

Previous studies using SEC-MALS and native gel electrophoresis have shown that wild type CtIP protein is a stable tetramer, and crystallographic studies provide a structural basis for this through a dimer-of-dimers arrangement of the N-terminal parallel coiled-coils (32). Disruption of this interface using the point mutation L27E is known to result in a dimeric form of CtIP (28, 32). Wild type CtIP and CtIP-L27E (with native masses of 404 kDa and 202 kDa respectively) run anomalously as widely dispersed smears on native gels (**Figure 2A**). Nevertheless, the dimeric L27E variant has a markedly greater mobility than the tetrameric wild type protein. The ΔC782 and ΔC608 deletion variants also ran as smears on these gels but with somewhat greater mobility than wild type as might be expected based on their reduced length. The band for CtIP-R100W appeared almost identical to wild type. Importantly, all three constructs ran with a significantly reduced mobility compared to the L27E dimer variant. These data suggest that ΔC782, ΔC608 and CtIP-R100W retain their wild type oligomeric state and confirms that they are all at least partially folded. To measure the molecular weight of the variant proteins we turned to SEC-MALS analysis (**Figure 2B**). Wild type CtIP ran as a single symmetrical peak which returned an average mass measurement of 384 kDa across the peak (close to the theoretical mass for a tetramer of 404 kDa). The disease variants also ran predominantly as single peaks with masses of 338, 262 and 376 for the ΔC782, ΔC608 and R100W proteins respectively. These are close to the expected values of 352, 272 and 404 for tetramers in all cases (compare the experimental mass data points with the expected mass dotted lines on **Figure 2B**).

**Figure 2.**
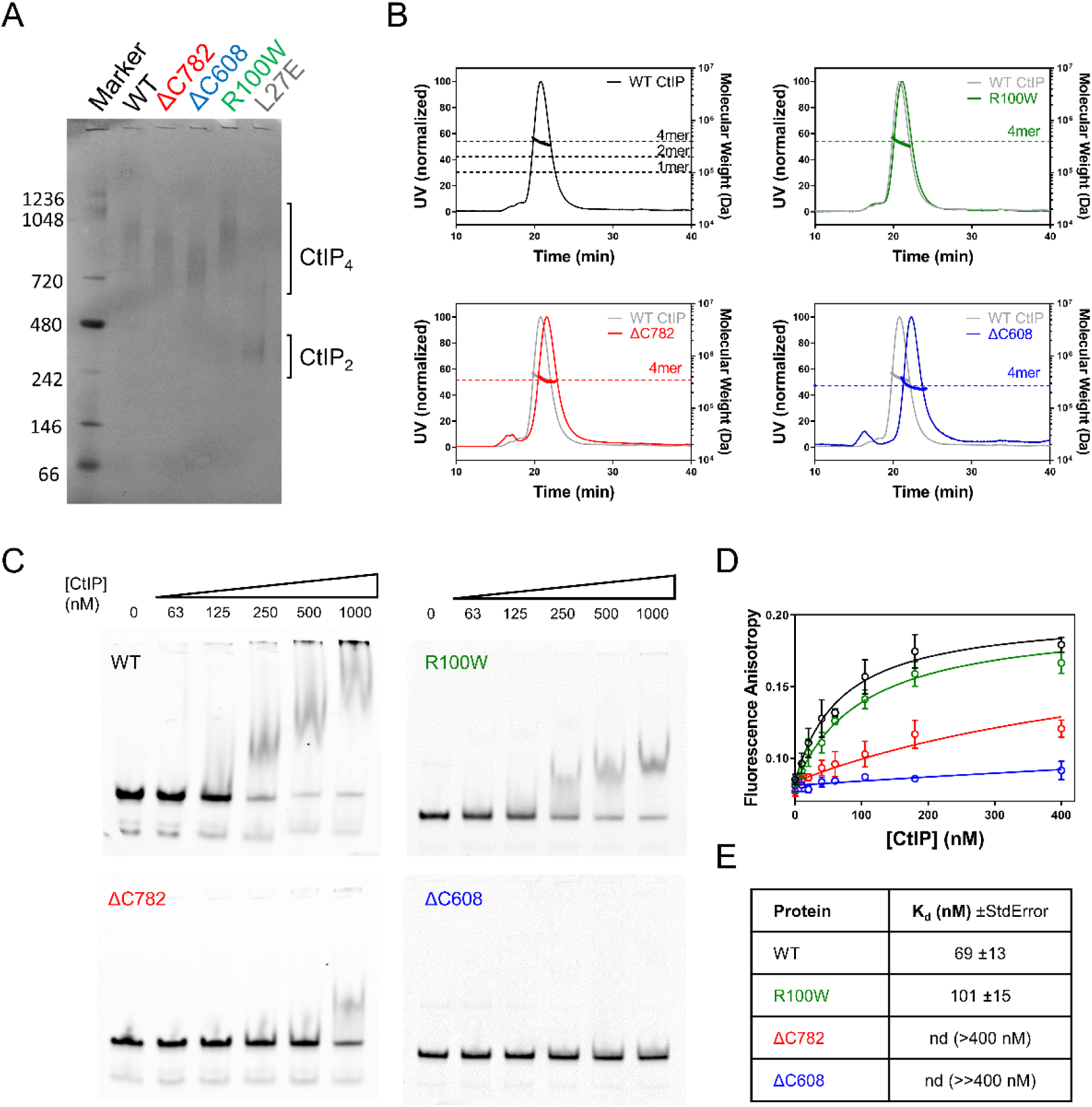
The CtIP disease variants tetramerise but cannot bind to DNA efficiently. (A) Native blue PAGE analysis of disease variant CtIP proteins (ΔC782, ΔC608 and R100W) in comparison to wild type and CtIP-L27E reference proteins which are known to form tetramers and dimers respectively. Note that each variant runs similarly to wild type, with a larger apparent mass than the dimeric L27E mutant protein. (B) SEC-MALS analysis of the same set of variant proteins, each compared to wild type CtIP. The dotted horizontal lines on each graph show the expected masses for the indicated oligomeric states of the protein of interest. Note that all the proteins run as tetramers. (C) Representative EMSA assays for wild type and variant CtIP proteins binding to a forked DNA substrate at the indicated (monomer) concentrations. Note that binding by the ΔC782 and ΔC608 proteins is severely compromised. (D) DNA binding isotherms for CtIP disease variants obtained by fluorescence anisotropy with a Cy5-labelled forked DNA molecule. The R100W variant displays slightly weaker binding than wild type whereas the ΔC782 and ΔC608 are more severely compromised in agreement with the EMSA assays. (E) The data shown in panel D were fit to yield the Kd values shown (see Methods for details).

To analyse DNA binding by CtIP disease variants we employed electrophoretic mobility shift assays (EMSA) and fluorescence anisotropy. We used a forked DNA substrate (containing a 25 bp duplex with 20 base 3ʹ- and 5ʹ-ssDNA overhangs) as we had established previously that this binds tightly to CtIP (28). In EMSA, wild type CtIP completely shifted the labelled DNA molecule yielding poorly defined bands within the gel and the wells as observed previously ((28); **Figure 2C)**. The CtIP-R100W variant also bound to the DNA with slightly higher concentrations of protein required to achieve a similar degree of shifting. In contrast, DNA binding was detectable but severely impaired in the ΔC782 variant and was undetectable at the highest concentration tested in the ΔC608 variant. The same forked DNA substrate was used in quantitative fluorescence anisotropy assays to produce binding curves for the wild type and variant CtIP proteins (**Figure 2D and 2E**). The wild type and CtIP-R100W data were well fit to a simple binding model to yield dissociation constants (Kd) of 66 and 115 nM respectively (see Methods for details). Binding was too weak to determine reliable Kd values for either of the deletion variants. However, ΔC782 did show evidence for partial binding at the highest concentration tested (400 nM) whereas ΔC608 did not. The two DNA binding assays are in broad agreement and show that R100W retains the ability to bind DNA albeit with a ∼2-fold reduced affinity compared to wild type, whereas both ΔC deletion variants are more severely compromised in their ability to interact with DNA.

### The CtIP ΔN variants bind to DNA but cannot tetramerise

We next performed analogous experiments using the ΔN variants (**Figure 3**). The ΔN500 protein was purified using SEC and reproducibly eluted as two peaks outside of the void volume of the column (**Supplementary Figure 1A**). These samples were stored separately as ΔN500a and ΔN500b. When the ΔN500a preparation was re-run on a SEC-MALS column, we observed the same two peaks (albeit not in the same volume ratio) implying an equilibrium between two oligomeric states that is established slowly. MALS analysis of these two peaks showed that they represent monomeric and dimeric populations (**Figure 3A**). The ΔN541 construct behaved in a similar way, running as both a monomeric and dimeric peak during SEC purification and SEC-MALS analysis (**Figure 3A**). In contrast, the ΔN790 construct ran as a single peak in SEC but produced equivocal SEC-MALS measurements in which the apparent mass varied significantly across the peak (ranging from 1.5-3.1 times the expected monomeric mass; **Supplementary Figure 1B**). Therefore, we also performed native mass spectrometry with this variant which showed that it was monomeric (**Figure 3B**). Interestingly, the experimental molecular mass for the ΔN790 construct (15421.50 ± 0.11 Da) was 63 Da greater than the expected mass for the monomeric polypeptide alone. This value is consistent with the binding of a single zinc atom to each ΔN790 monomer which may be co-ordinated by a C3H1 motif formed by conserved residues C813, C816, C835 and H838 (38), although this observation requires further experimental scrutiny.

**Figure 3.**
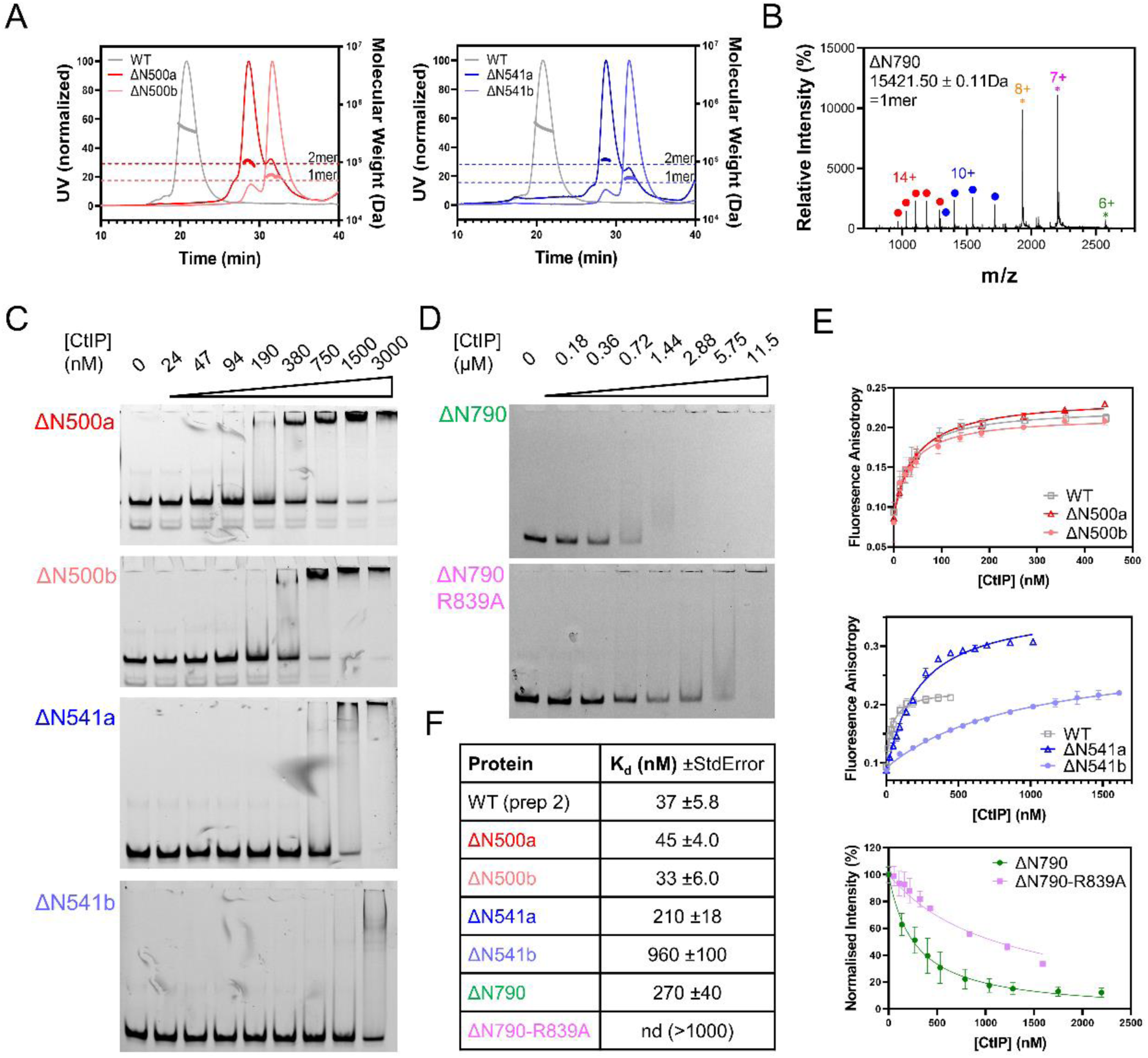
The CtIP ΔN variants bind to DNA but cannot tetramerise. (A) SEC-MALS analysis of the ΔN500/541 proteins. The ΔN500 protein is resolved as two peaks on SEC (which we call a and b, see main text). These were then individually re-run on a SEC column coupled to MALS analysis. Thin lines show UV absorbance whereas thick lines show the molecular weight calculation from the MALS signal. Peak a (red) is dimeric whereas peak b (pink) is monomeric. The grey trace shows data for WT CtIP for comparison. Similarly, the ΔN541 protein is also resolved as two peaks on SEC (a and b) which are dimeric (dark blue) and monomeric (light blue) respectively. (B) Native mass spectrum for the ΔN790 variant shows a monomer. The measured mass of the construct (15421.50 ± 0.11) is slightly greater than the theoretical mass (15358.56 Da) and consistent with the presence of a single bound Zn^2+^ atom. Charge states are colour-coded with green, magenta and orange asterisks representing folded low-charge states and with red or blue asterisks representing partially folded or unfolded high-charge states. (C) EMSA assays showing DNA binding by the ΔN500/541 variants. All variants bind DNA although affinity is reduced to differing extents (see main text for discussion). The CtIP (monomer) concentrations used are indicated above the gels. (D) EMSA assays showing DNA binding by the ΔN790 variant. The CtIP (monomer) concentrations used are indicated above the gels. (E) DNA binding isotherms based on fluorescence anisotropy (ΔN500/ΔN541) or quenching (ΔN790). Error bars represent the standard error of three experiments. (F) The data shown in panel E were fit to yield Kd for the interaction (see Methods for details).

In EMSA experiments (**Figure 3C**), the ΔN500 variant shifted the DNA substrate at similar concentrations to wild type but there was a qualitative change in the pattern of bands observed. Unlike wild type CtIP, ΔN500-DNA complexes formed discrete complexes within the gel, implying that removal of the coiled-coils results in a more conformationally homogenous system. EMSA gels for the ΔN500 variant were very similar regardless of whether material from SEC peak a (dimer) or b (monomer) was used in the experiments. Binding of ΔN541 to DNA was weaker than WT and ΔN500, but the shifted bands again appeared more discrete than the smeared bands that are always observed for wild type. The ΔN541a sample displayed a higher affinity for DNA than ΔN541b, possibly suggesting that a dimeric form of ΔN541 binds more tightly to DNA than a monomer. Finally, the ΔN790 variant shifted the forked substrate into the wells of the gel but with a reduced affinity compared to wild type (**Figure 3D**).

Fluorescence anisotropy measurements were in broad agreement with the gel shifts (**Figures 3E and 3F**). These showed that ΔN500 retained wild type DNA binding affinity, displaying Kd values of 45 nM and 33 nM for material from peak a and b respectively. Note that the wild type measurement here (Kd = 37 nM) represents an independent biological repeat of that used in comparison with the ΔC variants above. On this basis, the affinity of the ΔN500 variant for DNA is the same within error as wild type. The ΔN541a and ΔN541b variants bound to DNA less tightly than CtIP-WT with Kd values of 210 and 960 nM, respectively. Interaction of ΔN790 with Cy5-fork DNA resulted in a large fluorescence quenching, which is also apparent in the EMSA gels, implying that its mode of binding is qualitatively different to other variants. Binding isotherms were generated by exploiting this fluorescence quenching and yielded a Kd value of 270 nM. Since the ΔN790 construct behaved differently in fluorescence-based binding assays compared to wild type and other variants, we performed additional site-directed mutagenesis experiments to confirm the observed binding was a property of the CtIP deletion mutant itself rather than a contaminant. The ΔN790 protein retains a conserved arginine residue (R839) that has been shown to be important for DNA binding in wild type CtIP (28, 34, 35). Substitution of this arginine to alanine within the ΔN790 variant reduced its affinity for DNA measured either by EMSA or fluorescence anisotropy (**Figures 3E and 3F**).

### Only full length CtIP promotes efficient bridging of distant DNA segments

In a previous work, we used Atomic Force Microscopy (AFM) to directly observe the bridging of DNA molecules by CtIP and found that dephosphorylated CtIP (λ-CtIP) was more effective in bridging DNA molecules than untreated WT-CtIP (28). However, this imaging-based assay was semi-quantitative and we were unable to provide information on the stability and kinetics of CtIP-mediated bridging. To better characterise and compare the DNA bridging ability of the wild type and different CtIP variants, we used magnetic tweezers to study protein-mediated interactions between distant DNA segments by cycling between an extended and relaxed state of the DNA and monitoring reductions in the extension of the molecules due to protein bridging. For these experiments, we fabricated DNA molecules of ∼4.3 kbp with two modified ends, which were selectively attached to a glass surface and a superparamagnetic bead. This arrangement enables the application of force to the DNA molecule using a pair of magnets (**Figure 4A**). To test for DNA-bridging by CtIP, we cycle between high (4 pN, 3 min) and low (0.1 pN, 1 min) force regimes. Bridging events occurring during the low-force phase are revealed when the application of high force fails to return the substrate to its full extended conformation (Δz, reduction in DNA extension, **Figure 4A)**. This provides the basis for a single-molecule DNA bridging assay which can quantify DNA bridging efficiency and measure the kinetics of un-bridging.

**Figure 4.**
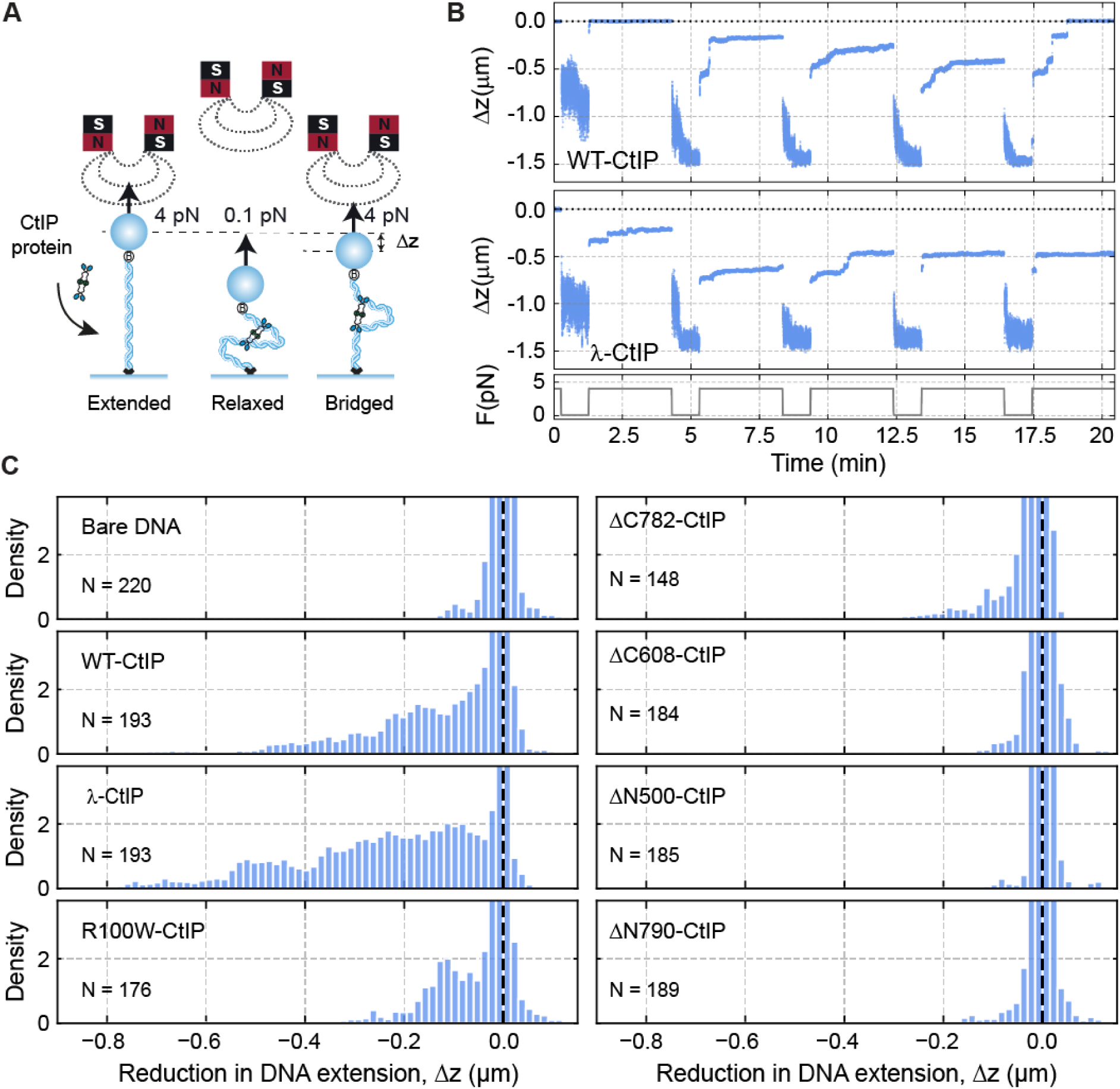
Only full length CtIP can promote bridging of distant DNA segments. (**A**) Cartoon of the magnetic tweezers assay to study dsDNA bridging. An intramolecular interaction results in the shortening (Δz) of the tether. (**B**) Representative time courses of two DNA molecules iteratively pulled in the presence of wild-type CtIP (WT-CtIP, top panel) and dephosphorylated CtIP (λ-CtIP, bottom panel). (**C**) Histogram of the relative reductions (Δz) for DNA molecules in absence (N = 220) and in the presence of WT-CtIP (N = 193), λ-CtIP (N = 193), R100W CtIP (N = 176), ΔC782 CtIP (N = 148), ΔC608 CtIP (N = 184), ΔN500 CtIP (N = 185) and ΔN790 CtIP (N = 189). The peak at Δz = 0 μm represents all extended states and the densities for Δz < 0 μm represent all bridged states through CtIP interactions.

Both WT-CtIP and λ-CtIP effectively promoted the interaction between distant DNA segments of a tethered DNA (**Figure 4B**). Raw data obtained from magnetic tweezers experiments are time-courses of DNA extension while cycling between high and low applied force. In these representative examples, we observed that, in some high-force steps of each cycle, the tether could not reach the nominal extension of the DNA determined in the absence of protein (dotted line in **Figure 4B**, see **Supplementary Figure 2** for other representative time courses). We interpreted this as the intramolecular bridging of two parts of the same DNA molecule mediated by CtIP. Interestingly, the high-force steps often displayed a continuous increase of the extension over time, which eventually reaches a stable value within the 3-min step. This can be interpreted as originating from the continuous breakage of multiple DNA-bridged interactions. A more detailed classification of the types of interaction events we observed and associated analyses are given below.

Our assay allowed us to follow tens of beads simultaneously and to perform a statistical analysis of Δz measured in the high-force step over all the molecules (**Figure 4C**). Data for WT- and λ-CtIP showed a clear tail in the extension distribution consistent with the reductions in length observed in individual time-courses (**Figure 4B and 4C**, see **Supplementary Figure 2** for other representative time courses). Interestingly, λ-CtIP showed a higher capacity to interact with distant remote sites, with some length reductions bigger than 1 μm. Experiments using CtIP variants with impaired DNA binding (ΔC608) or tetramerization (ΔN500/ΔN790) resulted in a signal comparable to the no protein control (**Figure 4C**). The disease-related variants ΔC782 and R100W showed impaired but nevertheless significant bridging which will be explored in greater detail below. Together, these data indicate that CtIP can bridge distant parts of the same DNA molecule and that efficient bridging requires both the DNA binding and tetramerization activities of CtIP.

We classified the different types of interactions observed in multiple time-courses for each of the different proteins, defining five types of events for each force cycle in order of increasing bridging stability. Class I events are those in which the DNA returns immediately to its full extension and there is no interaction (**Figure 5A-I**). Class II are those in which a bridging event is initially observed but this ruptures within the high-force step to recover the original DNA extension (**Figure 5A-II**). These events likely represent individual bridging interactions mediated by CtIP with a lifetime defined by τ and are analysed in greater detail below. Class III are bridging events with partial ruptures, where the full extension is not recovered within the high-force step, indicating the presence of multiple bridging events on the same DNA tether (**Figure 5A-III**). Class IV represents bridging events that do not rupture at all during the high force step (**Figure 5A-IV**). Finally, Class V are those exceptionally stable events where bridging is retained for more than one force cycle but for less than the entire experimental timecourse (five cycles; **Figure 5A-V**).

**Figure 5.**
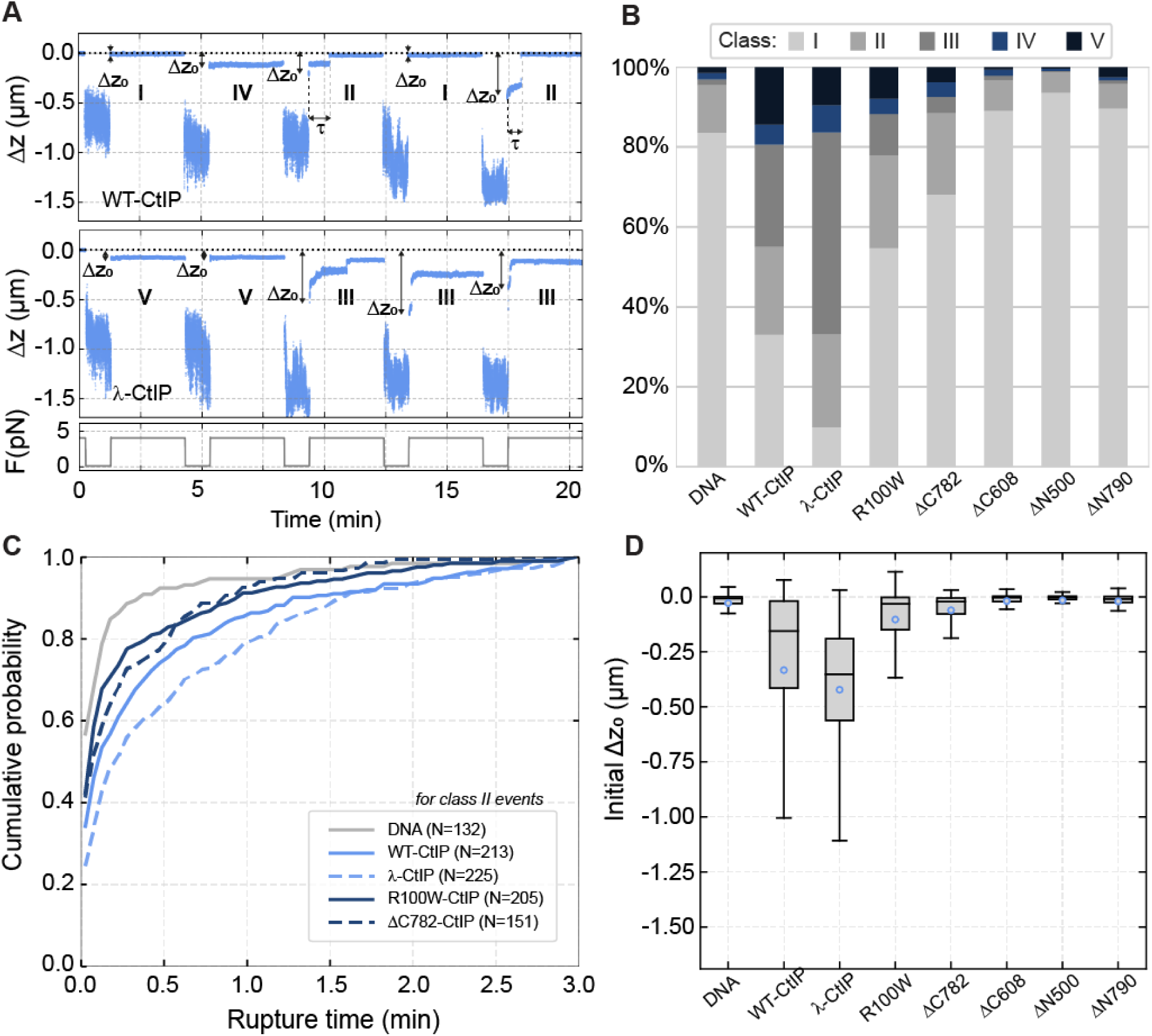
Analysis of DNA-bridging events by CtIP and CtIP variants. (**A**) Individual time courses with representative Class I-V events. Class I events (**I**) represent no observed bridging. Class II (**II**) represent single bridging events which rupture and return to full extension and where τ indicates the lifetime. Class III (**III**) are those events in which an initial bridge undergoes one or more ruptures but the DNA fails to return to full extension. Class IV (**IV**) and Class V (**V**) both represent highly stable bridging events over the entire high force step. For Class V events, the bridging remains stable over multiple pulling cycles. ΔZ_0_ represents the initial DNA extension at the beginning of each high force step. (**B**) Frequency of |Class I-V events for the different CtIP variants. (**C**) Cumulative probability of a rupture event for bare DNA, WT-CtIP, λ CtIP, R100W CtIP and ΔC782 CtIP reflecting the propensity of the DNA to un-bridge for each condition tested. (**D**) Boxplot representation of the initial DNA length reductions (ΔZ_0_) obtained for the different CtIP variants. Small empty circles indicate the average reduction. Boxes covers the inter-quartile range (IQR, Q1 to Q3), horizontal lines mark the median and upper/lower whiskers indicate Q1-1.5IQR and Q3+1.5IQR, respectively.

WT-CtIP and λ-CtIP interactions were mostly classified as Class II and III (**Figure 5B**). However, partial ruptures (Class III) were much more frequent for λ-CtIP. This probably reflects the presence of multiple proteins simultaneously bridging sections of the same DNA, such that breakage of a fraction of the tethers does not result in the full extension of the DNA and is consistent with a higher affinity observed between λ-CtIP and DNA (compared to wild type) (28). The average lifetime of the Class II interaction was 26.3 s for WT-CtIP and 33.9 s for λ-CtIP (**Supplementary Figure 3**). From the lifetimes of all the events, we determined the cumulative probability of a rupture as a function of time. In the absence of protein, most of the molecules are fully extended (90% probability) after 30 s of pulling, whereas in the presence of WT-CtIP there is a 75% probability reducing to 65% for λ-CtIP (**Figure 5C**). Therefore λ-CtIP is not only more efficient in binding and forming the intermolecular tethers between DNA segments, but also forms bridges which are subsequently more resistant to breakage compared to WT-CtIP.

As the bridging of DNA segments occurs during the low-force step, the initial extension for the first time-points of each high force step reveals whether (or not) successful bridging initially occurs, defining an event as Class II or greater (**Figure 5A**). Interestingly, the DC782 and R100W variants presented a fraction of Class II events that was comparable to WT- and λ-CtIP (**Figure 5B**). However, a detailed analysis of the initial Δz_0_ value for ΔC782 revealed mainly short-range bridges which break prematurely (average lifetime = 17.5 s, **Figures 5C-D** and **Supplementary Figure 3**)). Thus, although bridging of DNA by ΔC782 is possible, it is highly impaired in comparison to WT-CtIP and more susceptible to the application of force. R100W bridges somewhat larger segments of DNA but also displays a shorter average lifetime (18 s; **Figures 5C-D** and **Supplementary Figure 3**). The N-terminal deletion variants failed to support bridging, displaying mostly Class I events comparable to the no protein control (**Figures 5B-D** and **Supplementary Figure 3**).

Because CtIP is involved in the repair of DSBs, we next asked whether CtIP could efficiently tether broken DNA ends. We fabricated substrates containing DNA branches terminating in forked DNA ends and repeated the bridging experiments as before (**Supplementary Figure 4A**). We again observed CtIP-dependent DNA bridging, but there was no evidence of a prominent Δz peak at the expected distance of -0.2 μm that would indicate a specific end-to-end bridging of the fork ends (**Supplementary Figure 4B-C**, see legend for further details). This observation is consistent with our analyses above and with previous work which suggested that CtIP has no special affinity for free DNA ends *per se* but may be retained better on DNA containing blocked ends consistent with DNA sliding (28).

### Non-specific CtIP-DNA interactions underpin rapid and processive 1D diffusion

To characterise the binding of CtIP to DNA directly, we imaged the protein interacting with a long duplex molecule using a combination of optical tweezers and confocal fluorescence microscopy (C-Trap, Lumicks) (39). For these experiments, we engineered a long DNA molecule (∼25 kbp) with biotin-labelled ends. This design enabled the association of two streptavidin-coated micron-size beads, each held within its respective optical trap (**Figure 6A**). After conducting controls to confirm the expected length and integrity of the DNA, we moved the molecule, which was held at a constant force of 15 pN, into a channel containing 10 nM AlexaFluor^TM^ 635-labelled CtIP. Labelling of CtIP was achieved by incubating biotin-labelled CtIP (Bio-CtIP) with the Streptavidin AlexaFluor^TM^ 635 conjugate (see the Methods for details). We then recorded the fluorescence of a region of interest that included the tethered DNA and both beads (**Figure 6B**). We also conducted confocal line scans between the beads at a rate of 22 ms per line. These line scans enabled us to generate kymographs, showing the direct binding and movement of CtIP on the DNA (**Figure 6C**). CtIP exhibited non-specific binding throughout the DNA molecule and was so efficient that full DNA coverage with 10 nM CtIP was achieved within one minute. Therefore, to capture individual protein dynamics, we restricted the incubation time to a few seconds and quickly moved the tethered DNA to a channel devoid of protein. In addition to limiting CtIP binding, this procedure also reduced the fluorescence background in the confocal images, allowing us to observe the behaviour of single proteins, as depicted in **Figure 6D**. Under these conditions, kymographs showed that individual proteins explored the entire length of the tether in a random manner (**Figure 6E**). Moreover, these excursions of CtIP on the DNA typically persisted for several minutes, reflecting a stable interaction. Analysis of individual trajectories following published procedures (40-42) allowed us to determine the diffusion constant of CtIP, D=3.9±0.2 µm^2^ s^-1^ (mean±SEM) (**Figure 6F**, and **Supplementary Figure 5**). Notably, this diffusion constant is relatively high, compared to other proteins analysed using the same methodology, e.g. ParB (10-fold), SMC5/6 (4-40-fold), or BRCA2 (100-fold) (42-44). This likely reflects a soft interaction with the DNA, likely through the backbone. These experiments provide further direct evidence of the efficient non-specific binding of CtIP to DNA and show that CtIP particles exhibit random and highly-processive 1D diffusion suggesting a potential mechanism for bridging distant parts of the DNA.

**Figure 6.**
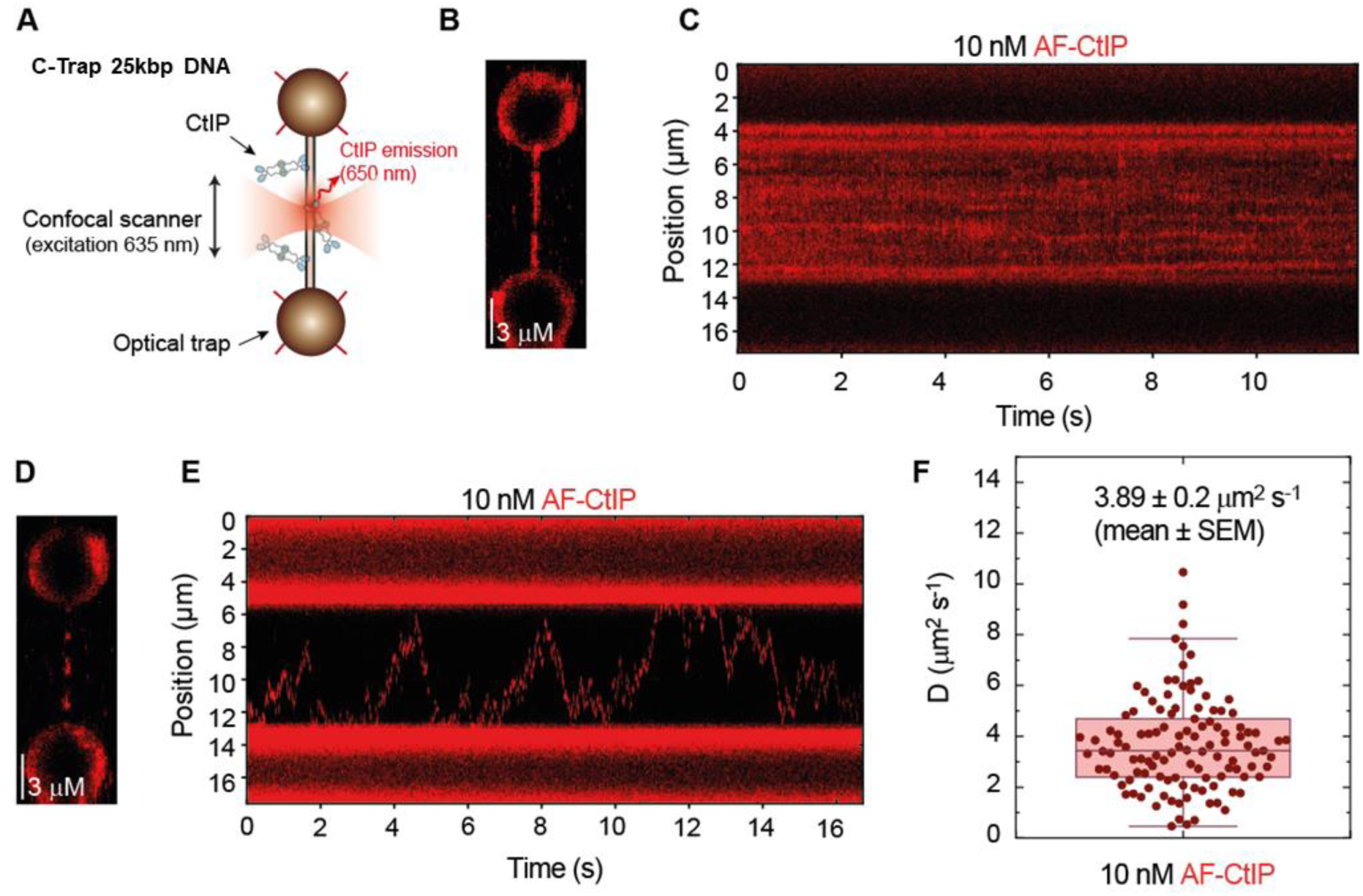
Non-specific CtIP-DNA interactions underpin rapid and processive 1D diffusion. **(A)** Cartoon of the experiment in which a dsDNA molecule is immobilized between two optically-trapped beads and held at 15 pN. Fluorescently-labelled CtIP was detected using a confocal microscope coupled to the dual optical tweezers. (**B**) Confocal scan of a DNA molecule densely covered by CtIP proteins after one-minute incubation with 10 nM CtIP. (**C**) Kymograph of CtIP non-specifically bound to dsDNA. (**D**) Confocal scan of a DNA molecule sparsely covered by CtIP proteins after ten-second incubation with 10 nM CtIP. (**E**) Kymograph of individual CtIP proteins diffusing along DNA. (**F**) Diffusion constants of CtIP obtained from individual CtIP trajectories.

## Discussion

In this work, we investigated the structure:function relationships which govern the oligomerisation, DNA binding and DNA bridging activities of CtIP; a protein which plays a key role in initiating DSB repair by homologous recombination. By comparing proteins with point mutations or which lack different N- and C-terminal regions of the protein we find that the C-terminal portion of the protein is responsible for DNA binding (**Figure 7**). A construct with the final 397 amino acids of the protein (ΔN500) has the same DNA binding affinity as wild type, and a construct retaining just 107 amino acids (ΔN790) also binds DNA albeit with moderately (∼4-fold) reduced affinity. This view is corroborated by CtIP variants containing C-terminal deletions which mimic variants of CtIP found in Seckel-2 and Jawad disease. Removal of the final 125 amino acids dramatically reduces DNA binding, and removal of the final 289 residues eliminates binding at the limit of sensitivity in our assays. Together, these data suggest that a major DNA binding domain resides in the C-terminal Sae2-like domain. Somewhat unexpectedly, we found that the ΔN790 variant (essentially just the Sae2-like domain) was monomeric (**Figure 7**). In previous work we showed that tetrameric CtIP-WT protein bound tightly to two forked DNA molecules and had interpreted that result as suggesting that the DNA binding domain(s) would dimerise to form a functional unit. However, although this work defines a minimal DNA binding domain in CtIP that can interact with DNA effectively as a monomer, it does not exclude the possibility that two such domains act together to form a continuous DNA binding surface in the context of the tetramer. Our observations are consistent with previous work which established that mutation of a conserved “RHR” motif in the Sae2-like domain reduces DNA binding affinity (28, 34, 35). Our DNA binding data also suggest that optimal interaction also requires amino acids residing in the region 500-782. In particular, the tighter binding displayed by ΔN500 compared to ΔN541 highlights a potential role for amino acids between 500-541. This region includes part of the “DR” motif, including residues K513 and K516, which have been implicated previously in DNA binding (36). Note that the ΔC608 variant, which includes the DR motif, is unable to bind DNA detectably in our assays. Therefore, although this region may contribute to the overall stability of the DNA interaction either directly or indirectly, it appears insufficient alone to facilitate tight binding. Primary structure analysis and current folding algorithms fail to provide confident insights into CtIP structure beyond the N-terminal coiled-coils (for which models are already available from crystallography) or very small regions of the C-terminus (45, 46). This probably reflects the extensive disorder that is predicted in C-terminal regions of this enigmatic protein. In this respect, we hope that the definition of a small minimal DNA binding locus associated with the small and conserved Sae2-like domain may provide a tractable target for hybrid structural approaches in the future.

**Figure 7.**
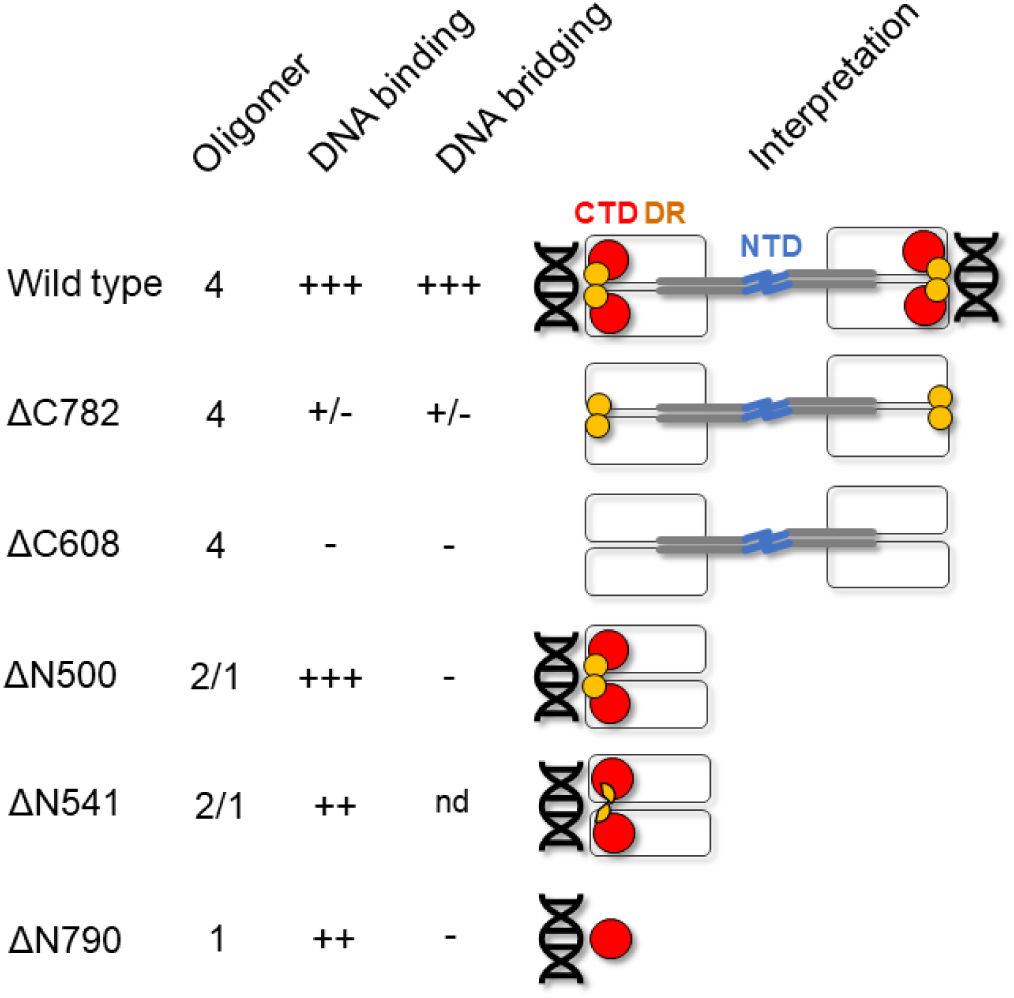
A model relating CtIP architecture to its DNA binding and bridging properties. Schematic summarising our results for wild type CtIP and variant proteins and their interpretation (see text for details). Colour coding is the same as in the Figure 1 primary structure diagram. Wild type CtIP (top row) is a tetrameric protein which can both bind DNA and efficiently bridge DNA molecules *in trans*. The protein adopts a dumbbell architecture where the N-terminal domains (NTD, blue) form a tetramerization interface at the centre of the dumbbell, with dimeric coiled coils (grey) extending away from the centre to the main body of the protein which forms a dimeric assembly consisting of the central disordered region (white) and the C-terminal Sae2-like domain (CTD, red). Single DNA molecules bind to each end of the dumbbell principally via the CTD and possibly supported by the DDR motif (yellow). The ΔC782 variant is equivalent to the variant produced in Seckel-2 disease and lacks the C-terminal Sae2-like domain. The protein is tetrameric, binds DNA very poorly, and consequently cannot bridge DNA effectively. Similarly, the ΔC608 variant, which is associated with Jawad syndrome and lacks both the CTD and the DDR motif, cannot bridge DNA due to its complete inability to bind DNA. The ΔN500 variant binds DNA as well as wild type. It cannot however bridge DNA molecules *in trans* as it lacks the NTD, cannot tetramerise, and instead exists as a mixture of dimeric and monomeric species. Similarly, the ΔN541 variant is unable to bridge DNA as it cannot tetramerise. Its DNA binding ability is somewhat compromised which may reflect the partial loss of the DDR motif. The ΔN790 variant, equivalent to the CTD domain alone, is proficient in DNA binding but cannot bridge DNA as it is monomeric. Nd = not determined.

Wild type CtIP not only binds DNA but can also efficiently bridge DNA segments *in trans*, a property that is likely to be crucial in its proposed role as a hub protein for initiating DNA repair at breaks. In this work we employed quantitative assay for CtIP-dependent bridging related to the molecular forceps assay that has previously been applied to non-homologous end joining factors (47-50). Only full length CtIP constructs were capable of efficient DNA bridging and did so without any strong preference for tethering ends (**Figure 7**). Dephosphorylated CtIP presented a higher efficiency of bridging consistent with tighter DNA binding affinity (28). CtIP-dependent DNA bridges were stable even at moderate opposing forces, an observation which lends credence to the idea that CtIP-dependent DNA tethering may help to orchestrate DNA break repair by maintaining the proximity of the participating molecules. As was to be expected, the ΔC variants (that bind to DNA poorly or not at all) did not bridge DNA molecules. These variants mimic the proteins produced in Seckel-2 and Jawad syndromes and this result is therefore consistent with the idea that defective DNA bridging may contribute to the disease phenotype. Importantly, we also found that the ΔN variants, which retain DNA binding at near wild type (ΔN500) or reduced affinities (ΔN541 and ΔN790), are incapable of DNA bridging. These results indicate that DNA binding is required but not sufficient for bridging and that the successful tethering of the DNA also requires tetramerization. The properties of the R100W mutant were especially interesting. This mutant, which is associated with Seckel-like disease and breast cancer, tetramerises normally and has a mild DNA binding defect, yet bridging is rather severely compromised. The mutation is in the coiled coil domains of CtIP, close to a conserved Zn^2+^ binding site, but nowhere near any of the proposed DNA binding loci (33). Therefore, the impact of this mutation on DNA bridging (and to a lesser extent binding) might be explained by allosteric effects associated with the structural integrity or dynamics of the coiled coils. The CtIP tetramer forms a particle with a dumbbell architecture that is held together by a dimer-of-dimers interface formed by the N-terminal coiled-coil domains. A simple interpretation of our data is that the distal ends of the dumbbell form the major DNA binding sites that are dimeric in nature but retain DNA binding affinity as monomers (**Figure 7**). It is possible that the relative orientation of the DNA binding domains or the flexibility of the intervening coiled coils are important for efficient bridging. The significance of CtIP-dependent DNA bridging for DNA end resection and strand exchange is yet to be fully explored. However, it is known that point mutations (L27E) which prevent tetramerization are non-functional in vivo. In the future, it will be interesting to determine whether DNA bridging is re-enforced by additional resectosome proteins such as MRN, whether it is maintained during processive DNA end resection, and whether bridging involves two broken DNA ends and/or additional *in trans* interactions with the sister chromosome which acts as the homology donor molecule in downstream recombination reactions.

## Materials and Methods

### Wild type and variant CtIP purification and labelling

Wild type (WT) CtIP cloned into a pACEBac1 vector (28) was used as a template to generate deletion variants for expression in insect cells using the QuickChange II XL site-directed mutagenesis kit (Agilent Technologies). For bacterial expression of the ΔN790 variant, a synthetic gene optimised for *E. coli* expression was purchased from Thermo Fisher (Gene Art) and cloned into the pET28a plasmid. Expression and purification of WT-CtIP and ΔC variants were carried out as detailed in (28). Purification of the ΔN500 and ΔN541 variants was performed with modifications to the same procedure. Briefly, Sf21 cells seeded at 500,000 cells/mL were transfected with recombinant bacmid DNA to generate the initial baculovirus (P1) in Insect-XPRESS^TM^ Protein-free Insect Cell Medium (Lonza). The P1 virus was amplified two times (P2 and P3) by infecting larger volumes of Sf21 cells. For large scale protein production, High-Five cells seeded at 2 million cells/mL were infected with P3 viruses and harvested after 72 h by centrifugation at 1000 g for 10 min. Cells were resuspended in buffer S containing 50 mM Tris pH 8, 50 mM NaCl, 0.5 mM TCEP, 10 % glycerol, supplemented with Roche cOmplete™ Protease Inhibitor Cocktail EDTA-free and lysed by sonication on ice for 3 minutes. The lysate was clarified by centrifugation at 50,000 g for 30 minutes at 4°C and applied to pre-equilibrated StrepTactin resin (IBA Life Sciences). The soluble extract was incubated with the resin for one hour at 4°C with constant rotation. The resin was washed five times in batch with 10 column volumes (CV) of buffer S and 2 CV of buffer W (50 mM Tris-Cl pH 8, 200 mM NaCl and 0.5 mM TCEP). The bound protein was eluted in 3 CV buffer W supplemented with 2.5 mM desthiobiotin (IBA Life Sciences). This material was then applied to a MonoQ 5/50 GL (Cytiva) column by diluting the salt concentration to approximately 100mM. The proteins were washed and eluted with a salt gradient up to 500 mM NaCl in buffer S. Protein fractions were pooled and concentrated to a volume less than 500 µL using centrifugal filters (Amicon) of appropriate molecular weight cut-offs and applied to a Superose 6 Increase 10/300 GL (Cytiva) in buffer G (20 mM Tris-Cl pH 8, 200 mM NaCl, 0.5 mM TCEP). Finally, the proteins were concentrated as required, supplemented with 10 % glycerol and flash frozen in liquid nitrogen and stored at -80 °C. The ΔN790 variant was expressed in *E. coli* using standard IPTG induction techniques and was purified in the same manner as the ΔN500 and ΔN541 variants with two exceptions; all buffers used were set to pH 7 and, following elution from StrepTactin resin, the sample was applied to a MonoS 5/50 GL column (Cytiva) instead of a MonoQ column.

To produce biotinylated CtIP (Bio-CtIP), a modified version of the WT-CtIP pACEBac1 plasmid was constructed containing a C-terminal Avi-tag upstream of the cleavable StrepII tag. This protein expressed and purified in an identical manner to wild type. After StrepII-tag cleavage and subsequent concentration to 50 µM, CtIP-Avi was treated with BirA (gift from Charles Grummit, University of Bristol) and biotin to site-specifically biotinylate the protein (51). After SEC purification (carried out as before) the protein was concentrated and then tested for biotinylation efficiency by PAGE gel (measured as ∼100%). Conjugation of the biotinylated CtIP protein to streptavidin did not affect its biochemical activities. Fluorescence labelling of CtIP was performed by incubation of Bio-CtIP with the Streptavidin AlexaFluor^TM^ 635 conjugate (ThermoFisher) in a 1:1 molar ratio for 10 minutes on ice (Bio-CtIP-AF635), and then diluted to a final concentration of 10 nM in 200 µl of reaction buffer (10 mM Tris-Cl pH 7.6, 20 mM NaCl, 1 mM DTT and 0.1 mg/mL BSA), supplemented with 10 µM biotin to neutralize the remaining streptavidin molecules.

### Preparation of DNA substrates for DNA binding assays

The fork DNA substrate used in DNA binding assays were prepared by annealing HPLC-purified synthetic oligonucleotides (Fork F and Fork R) with a 5’ Cy5 or HEX label on one strand (see **Supplementary Table 1** for sequences). The annealing reaction was performed by heating equimolar amounts of both strands (25 µM each) in 50 mM Tris-Cl pH 8, 150 mM NaCl, 1 mM EDTA for 10 minutes and allowing them to cool slowly overnight.

### Size exclusion chromatography coupled to multi-angle light scattering (SEC-MALS)

Protein samples within the concentration range 0.5-1 mg/mL were injected onto pre-equilibrated Superose 6 10/300 size-exclusion chromatography column (GE Healthcare) in buffer containing 20 mM Tris-Cl, pH 8.0, 200 mM NaCl, 0.5 mM TCEP. The DAWN HELEOS II MALS detector (Wyatt Technology) and an Optilab T-rEX differential refractometer (Wyatt Technology) were used to record the data. Analysis was performed using the ASTRA 6 software (Wyatt Technology). Graphs were plotted using GraphPad PRISM.

### Native nanoESI-MS

Mass spectra were recorded on a Synapt G1 HDMS mass spectrometer (Waters) set to an m/z range of 650-8000 and calibrated internally using caesium iodide. Both EcN4 and InN4 were dialysed into 20 mM ammonium acetate solution. 8 µL sample was pipetted into a borosilicate nano-emitter (made in-house) and loaded onto the nano electrospray ionisation (ESI) source. Instrument conditions were optimised for ion desolvation whilst minimising denaturing the samples; capillary voltage: 1.2 kV, sampling cone: 30 V, extractor cone: 0.3 V, trap collision energy: 6 V, transfer collision energy: 8 V, source temperature: 30 °C, backing pressure: 2.12 mbar, trap pressure: 1.11e-2 mbar, IMS pressure: 4.36e-4 mbar and TOF pressure 1.26e-6 mbar. Data acquisition and processing were performed in MassLynx 4.2 (Waters).

### Electrophoretic mobility shift assays

2.5 nM of 5’-Cy5 labelled fork was mixed with increasing amounts of CtIP and CtIP variants in buffer containing 20 mM Tris-Cl pH 8.0, 100 mM NaCl, 1 mM DTT, 0.1 mg/mL BSA, 5 % glycerol. The mixture was incubated at room temperature for a minimum 10 minutes before loading onto a 6 % native polyacrylamide gel. The samples were electrophoresed for 35 minutes at 150 V in 1X Tris-Borate-EDTA (TBE) buffer and the bands were visualised using a Typhoon scanner (GE Healthcare).

### Fluorescence-based DNA binding assays

Anisotropy measurements were performed by titrating increasing amounts of concentrated protein stocks into buffer containing 20 mM Tris–Cl (pH 8.0), 20 mM NaCl, 1 mM DTT and 5 nM 5’-HEX labelled fork. A Horiba Jobin Yvon FluoroMax fluorimeter was used to record the measurements with excitation and emission wavelengths of 530 and 550 nm respectively. One-minute incubations were performed before recording each measurement and an average of two readings was taken for each measurement. The binding curves were plotted and analysed as detailed in (28). For the ΔN790 variant, fluorescence intensity scans were performed and the change in intensity values at 550 nm were recorded and plotted to calculate the binding affinity.

### Magnetic Tweezers DNA bridging assay

#### DNA substrates

Magnetic tweezers experiments were performed on two types of DNA molecules: a ‘branched’ DNA, which features two short dsDNA fragments branching from the central dsDNA segment, and an ‘unbranched’ DNA, which lacked such branching. The fabrication of these dsDNA constructs has been described in detail in a previous work (50). Briefly, both DNA designs are composed of a central dsDNA segment of 4292 bp (See **Supplementary Table 2** for sequence) flanked by two labelled handles. One handle (1 kbp) is labelled with digoxigenin to ensure the DNA immobilization onto the lower surface of the flow cell, while the other one (140 bp) is biotinylated for the attachment of a streptavidin-covered superparamagnetic bead. The branched DNA design differentiates from the unbranched by two fork-terminated fragments, separated by 689 bp, protruding from the central dsDNA segment. Each fork-branch features two 20 nt-long polydTs linked to a 40 bp-long dsDNA fragment. See **Supplementary Table 1** for sequences of oligonucleotides.

#### Magnetic tweezers instrument

The magnetic tweezers equipment used in this work was similar to others described in the literature (52, 53). Briefly, the equipment consists of a pair of permanent magnets and a flow chamber arranged on top of an inverted microscope. To create the flow chamber, a sandwich comprising an upper coverslip with two small holes (serving as inlet and outlet), a double-parafilm spacer, and a lower coverslip covered with polystyrene is assembled and subsequently sealed through the application of heat. For the experiments, DNA molecules previously attached to superparamagnetic beads were flushed onto the flow chamber and immobilized on the chamber’s lower surface. The permanent magnets placed above the flow chamber allowed the manipulation of the tethered DNA molecules by attracting the superparamagnetic beads. The inverted microscope is equipped with a high magnification oil-immersion objective connected to a CCD camera that records the position of the bead at 120 Hz. The distance-dependent force applied on the DNA molecules was computed from the Brownian excursions of the molecules (52). The extension of the DNA molecules was determined by comparing images taken at different focal planes. In this setup, up to 40 molecules could be tracked simultaneously and forces up to 5 pN were applied to 1 µm diameter beads.

#### In-chamber DNA Immobilization

First, the flow chamber was incubated at 4°C overnight with 25 ng/µL Digoxigenin Antibody (Bio-Rad) that adsorbed onto the polystyrene-covered lower surface. When preparing for an experiment, 8 µL of DNA (1.4 nM) diluted 1:300 in TE (10 mM Tris pH 8.0, 1mM EDTA) was mixed with 20 µL of 1 µm-diameter magnetic beads (Dynabeads MyOne Streptavidin T1, Thermo Fisher Scientific) diluted 1:10 in PBS-BSA (PBS supplemented with 0.4 mg/mL BSA (New England Biolabs)). During this step, the biotinylated handle of the DNA constructs attached to the streptavidin-covered magnetic beads. After 10 minutes, the excess DNA in solution was removed by trapping the beads with a magnet, discarding all the volume and adding fresh PBS-BSA. After washing three times, the DNA-beads were resuspended in 70 µL and flushed into the flow cell. The handle labelled with digoxigenin interacted with the anti-digoxigenin on the lower coverslip and the DNA molecules stay immobilized. After 15 minutes, the excess beads were washed away by flushing PBS (around 1 mL).

#### DNA bridging assay

To study the internal bridging of dsDNA a sequence of force cycles was applied to the DNA molecules in the presence of the proteins of interest. The sequence was the following: a short starting step of 10 s at a 4 pN force, followed by 5 repetitions of a relaxation step at 0.1 pN for 1 minute and a pulling step at 4 pN for 3 minutes. To start an experiment, the DNA molecules were immobilized inside the flow chamber as described in the previous section. In the next step, negative rotations were applied to the tethered beads at a force of 4 pN to identify those that are attached to the surface by more than one DNA molecule. These multiple-DNA tethers were discarded during data processing. A full cycle of high/low force steps was first performed on bare DNA molecules to confirm there is not bridging interactions in the absence of protein. Then, the protein of interest was flushed into the flow chamber and let to stabilize for a minute. Finally, the force sequence was applied. All experiments were performed in 10 mM Tris-Cl pH 7.6, 20 mM NaCl, 1 mM DTT and 0.1 mg/mL BSA, with 4 nM monomeric concentration of the protein of interest. Experiments that did not show interactions, such as those of DC608, DN500 and DN790 were also performed at a higher concentration of 20 nM to discard any concentration-derived effect. Finally, 10 µM biotin (Sigma-Aldrich) is included in experiments with proteins that contain strep-tags, such as DN500 and DN790.

#### Data processing

Datasets obtained from the magnetic tweezers’ experiments are analysed using custom Python scripts. These datasets contain the absolute extension of each molecule with respect to time and force. The first step of the data processing is to transform this absolute extension (z) to a relative DNA extension (Δz). This step reduces the impact of the variability between absolute extension of the tethers (mainly arising from different anchoring points). The resulting Δz is the difference between the extension at each timepoint with respect to the averaged initial extended value. If Δz is 0 μm, the molecule maintains its initial extension. Whereas Δz < 0 μm indicates reductions in extension and Δz > 0 μm elongations. Data from multiple DNA molecules and several repetitions are merged and processed to obtain a global distribution of relative extensions. The distributions are normalised so that the area under the histogram is 1. The population size (number of DNA molecules) for each protein condition is marked on each distribution as an N in the lower left corner. For the detailed analysis and classification of the events, a few parameters are extracted from each of the five pulling steps for each time course, for all molecules and CtIP variations. These parameters are the initial extension, the final extension and the rupture time. The pulling steps last for 3 minutes, so the initial extension (time = 0 min) is defined as the averaged reduction in length measured for the first 12 points of the pulling cycle, Δz_0_, and the final extension is the averaged reduction in length measured for the last 12 points of the 3-min step, Δz_3-min_. The rupture point is defined as the moment a molecule recovers its original extension (-0.05 μm < Δz < 0.05 μm). The rupture time, or lifetime of the bridging event, τ, is the time since the pulling starts until the rupture point. These parameters are used to classify the events following a specific criterion. Class I events are those where the molecule is not bridged and the measured Δz_0_ is within an acceptance range for an extended molecule around Δz = 0 μm (-0.05 μm < Δz_0_ < 0.05 μm). Class II includes the events in which the molecule is initially bridged and suffers an eventual rupture before the end of the 3-min step. For Class II events, the measured Δz_0_ is below and the Δz_3-min_ is within the extended molecule range (Δz_0_ < -0.05 μm and -0.05 μm < Δz_3-min_ < 0.05 μm). Class III represents the events that rupture partially and never reach the extended state, so both Δz_0_ and Δz_3-min_ are below the extended range and the there is a significant difference between the two extensions (Δz_0_ < -0.05 μm, Δz_3-min_ < -0.05 μm and |Δz_3-min_ -Δz_0_ | > 0.05 μm). Finally, both Class IV and Class V integrate all bridging events whose extension is immutable for the entire 3-min step, so |Δz_3-min_ -Δz_0_ | < 0.05 μm, where Class V events persist for more than one consecutive cycle and Class IV are limited to one cycle.

### Optical Tweezers and Confocal Microscopy Assay

#### DNA substrates

C-Trap experiments were performed on dsDNA molecules of 25427 bp. The central part of the DNA construct was obtained by digestion of a large homemade plasmid with BssHII (New England Biolabs) produced following published protocols (54). Without further purification, the fragment was ligated to highly biotinylated handles of ∼1 kbp ending in MluI. Handles for C-trap constructs were prepared by PCR (see **Supplementary Table 1** for primers) including 200 µM final concentration of each dNTP (dGTP, dCTP, dATP), 140 µM dTTP and 66 µM Bio-16-dUTP (Roche) using the plasmid pSP73-JY0 as template (55) followed by digestion with the restriction enzyme MluI (New England Biolabs). Labelled handles were ligated with the central part with T4 DNA Ligase (New England Biolabs) for 15 h at 16°C followed by 1 h at 37°C before heat inactivation, in the presence of BssHII enzyme to avoid tandem (double-length) tethers. These handles were highly biotinylated to facilitate the capture of DNA molecules in C-Trap experiments. The sample was ready for use without further purification. DNAs were never exposed to intercalating dyes or UV radiation during their production and were stored at 4°C (see **Supplementary Table 2** for sequence).

#### DNA binding and single-protein tracking assay

Single-molecule experiments were carried out using a dual optical trap combined with confocal microscopy (C-Trap; Lumicks) (39) applying the same protocols and procedures as described in (42, 56). The buffer for trapping the beads and DNA was 20 mM HEPES pH 7.8, 100 mM KCl and 5 mM MgCl_2_. The length and integrity of the DNA molecules was tested by performing force-extension curves in reaction buffer. Then, the trapped DNA was moved to the protein channel containing 10 nM Bio-CtIP-AF635 for confocal imaging. Confocal experiments employed a 635 nm excitation laser for the visualization of the AF-635 fluorophore, with an emission filter of 650-750 nm. The confocal laser intensity at the sample was set to 1.92 µW. To extend the lifetime of the fluorophore, confocal imaging of single CtIP diffusion was performed in the presence of an oxygen scavenger system composed by 5 mM 3,4-Dihydroxybenzoic acid (PCA) and 100 nM-Protocatechuate 3,4-Dioxygenase (PCD). For the analysis of diffusion constants, DNA molecules containing more than two CtIP particles were discarded. Imaging channels were passivated for 30 minutes with BSA (0.1 % w/v in PBS) prior to the experiment to prevent unspecific interaction of the protein with the surface. Kymographs were generated by single line scans between the two beads using a pixel size of 100 nm and a pixel time of 0.1 ms, resulting in a typical time per line of 22.4 ms. Experiments were performed in constant-force mode at 15 pN.

#### Data processing

We use custom Python scripts to access, visualize and export confocal data from Bluelake (Lumicks) HDF5 files obtained from C-Trap experiments. The quantification of the individual trajectories of CtIP-AF635 was performed using a custom LabVIEW program that provides the position of individual CtIP proteins along the DNA for a given time (t) (42). The length of the time courses was restricted to 2.5 s to increase the statistical sample. The mean square displacement (MSD) was then computed for a given time interval (Δt) and the diffusion coefficient (D) calculated as described in (40-42). A total of 115 trajectories was used for the diffusion coefficient calculation.

## Acknowledgements and Funding Sources

We are grateful to Ian Taylor (The Crick Institute) for access to and assistance with SEC-MALS measurements. Work in the MSD laboratory was supported by the Wellcome Trust (100401/Z/12/Z) and the Biotechnology and Biological Sciences Research Council (BB/V001817/1). SLB was supported by the Wellcome Trust Dynamic Molecular Cell Biology PhD programme at the University of Bristol. Work in the FMH laboratory was supported by grants PID2020-112998GB-I00, funded by Ministerio de Ciencia e Innovación (MICINN)/Agencia Estatal de Investigación (AEI/10.13039/501100011033)_FEDER, EU, and co-funded by the European Regional Development Fund (ERDF); grants Y2018/BIO4747 and P2018/NMT4443, funded by the Autonomous Region of Madrid and co-funded by the European Social Fund (ESF) and the ERDF; grant EUREXCEL Ref. 951214, funded by CSIC. Work in the FS laboratory was supported by the Biotechnology and Biological Sciences Research Council (BB/V001698/1).

