## Supplementary Information File for "DNA binding and bridging by human CtIP in the healthy and diseased states"

Including links/DOI for data access, Supplementary Tables 1 and 2, and Supplementary Figures 1-5.

### Links/DOI for data access

Part 1/3 - https://data.mendeley.com/preview/vdphmchspb?a=ddbdfe1f-92c9-412e-98bc-2f91e3106fbd

DOI:10.17632/vdphmchspb.1

Lokanathan Balaji, Shreya (2023), "Lokanathan-Balaji et al. bioRxiv 2023 (1/3)", Mendeley Data, V1, doi: 10.17632/vdphmchspb.1

Part 2/3 - https://data.mendeley.com/preview/kv7b4jkmt9?a=a761994b-bf26-459c-9d96-a51fcb75fae6

DOI:10.17632/kv7b4jkmt9.1

Lokanathan Balaji, Shreya (2023), "Lokanathan-Balaji et al. bioRxiv 2023 (2/3)", Mendeley Data, V1, doi: 10.17632/kv7b4jkmt9.1

Part 3/3 - https://data.mendeley.com/preview/2bnwfnr5jg?a=a019127f-a80b-4916-83ac-0f9cd46e3151

DOI:10.17632/2bnwfnr5jg.1

Lokanathan Balaji, Shreya (2023), "Lokanathan-Balaji et al. bioRxiv 2023 (3/3) - supplementary", Mendeley Data, V1, doi: 10.17632/2bnwfnr5jg.1

### Supplementary Table 1 – Sequences of oligonucleotides used in this study

| Fragment | Oligonucleotide | Sequence |
|---|---|---|
| MT DIG handle | 57.FMH_F2_BamHI-ApaI | GCGTAAGTGGATCCGGGCCCCGACTCACTATAGGGAGACCGGC |
| | JOE_R1 | AGTAAGCGCCGTCAGACCAG |
| MT BIO handle | FMH_F2_BsrGI | GCGTAAGTTGTACACGACTCACTATAGGGAGACCGGC |
| | 209.BsrGI 71short handle | CGATAACCAACTGGCGATG |
| C-Trap BIO handle | 207.FMH_F2_MluI | GCGTAAGTACGCGTCGACTCACTATAGGGAGACCGGC |
| | JOE_R1 | AGTAAGCGCCGTCAGACCAG |
| Forked-end branch | 94.P-Fork 20T EcoRV NotI | (Pho)TCAGCTCATGTCATCCTCAGCACACTTGACCGGCTAGGCAGGATATCATGCATGCGGCCGCCATGGAGGGTTTTTTTTTTTTTTTTTTTTT |
| | 95.Fork 20T EcoRV NotI | TTTTTTTTTTTTTTTTTTTTCCCTCCCATGGCGGCCGCATGCATGATATCCTGCCTAGCCCTCAGCTCAGCTAGCCTCAGCCTACAATCACC |
| Cy5/HEX Fork DNA | Fork F | [Cy5]/[HEX]GCTTGCTAGGACGGATCGCTCGAGGTTTTTTTTTTTTTTTTTTTTT |
| | Fork R | TTTTTTTTTTTTTTTTTTTTCCTCGAGCGATCCGTCCTAGCAAGC |

### Supplementary Table 2. dsDNA constructs used in this study

- Underlined: 63 nt-gaps created after digestion with the nicking enzyme Nt.BbvCI followed by denaturation.

| Fragment | Sequence |
|---|---|
| MT central fragment<br><br>(4292 bp) | GTACAGAAAACTCACGTTAAGGGATTTTGGTCATGAGATTATCAAAAAGGATCTTCACCTAGATCCTTTTAAATTAAAAATGAAGTTTTAAATCA<br>ATCTAAAGTATATATGAGTAAACTTGGTCTGACAGTTACCAATGCTTAATCAGTGAGGCACCTATCTCAGCGATCTGTCTATTTCGTTCATCCAT<br>AGTTGCCTGACTCCCCGTCGTGTAGATAACTACGATACGGGAGGGCTTACCATCTGGCCCCAGTGCTGCAATGATACCGCGAGACCCACGCTCAC<br>CGGCTCCAGATTTATCAGCAATAAACCAGCCAGCCGGAAGGGCCGAGCGCAGAAGTGGTCCTGCAACTTTATCCGCCTCCATCCAGTCTATTAAT<br>TGTTGCCGGGAAGCTAGAGTAAGTAGTTCGCCAGTTAATAGTTTGCGCAACGTTGTTGCCATTGCTACAGGCATCGTGGTGTCACGCTCGTCGTT<br>TGGTATGGCTTCATTCAGCTCCGGTTCCCAACGATCAAGGCGAGTTACATGATCCCCCATGTTGTGCAAAAAAGCGGTTAGCTCCTTCGGTCCTC<br>CGATCGTTGTCAGAAGTAAGTTGGCCGCAGTGTTATCACTCATGGTTATGGCAGCACTGCATAATTCTCTTACTGTCATGCCATCCGTAAGATGC<br>TTTTCTGTGACTGGTGAGTACTCAACCAAGTCATTCTGAGAATAGTGTATGCGGCGACCGAGTTGCTCTTGCCCGGCGTCAATACGGGATAATAC<br>CGCGCCACATAGCAGAACTTTAAAAGTGCTCATCATTGGAAAACGTTCTTCGGGGCGAAAACTCTCAAGGATCTTACCGCTGTTGAGATCCAGTT<br>CGATGTAACCCACTCGTGCACCCAACTGATCTTCAGCATCTTTTACTTTCACCAGCGTTTCTGGGTGAGCAAAAACAGGAAGGCAAAATGCCGCA<br>AAAAAGGGAATAAGGGCGACACGGAAATGTTGAATACTCATACTCTTCCTTTTTCAATATTATTGAAGCATTTATCAGGGTTATTGTCTCATGAG<br>CGGATACATATTTGAATGTATTTAGAAAAATAAACAAATAGGGGTTCCGCGCACATTTCCCCGAAAAGTGCCACCTAAATTGTAAGCGTTAATAT<br>TTTGTTAAAATTCGCGTTAAATTTTTGTTAAATCAGCTCATTTTTTAACCAATAGGCCGAAATCGGCAAAATCCCTTATAAATCAAAAGAATAGA<br>CCGAGATAGGGTTGAGTGTTGTTCCAGTTTGGAACAAGAGTCCACTATTAAAGAACGTGGACTCCAACGTCAAAGGGCGAAAAACCGTCTATCAG<br>GGCGATGGCCCACTACGTGAACCATCACCCTAATCAAGTTTTTTGGGGTCGAGGTGCCGTAAAGCACTAAATCGGAACCCTAAAGGGAGCCCCCG<br>ATTTAGAGCTTGACGGGGAAAGCCGGCGAACGTGGCGAGAAAGGAAGGGAAGAAAGCGAAAGGAGCGGGCGCTAGGGCGCTGGCAAGTGTAGCGG<br>TCACGCTGCGCGTAACCACCACACCCGCCGCGCTTAATGCGCCGCTACAGGGCGCGTCCCATTCGCCATTCAGGCTGCGCAACTGTTGGGAAGGG<br>CGATCGGTGCGGGCCTCTTCGCTATTACGCCAGCTGGCGAAAGGGGGATGTGCTGCAAGGCGATTAAGTTGGGTAACGCCAGGGTTTTCCCAGTC<br>ACGACGTTGTAAAACGACGGCCAGTGAGCGCGCGTAATACGACTCACTATAGGGCGAATTGGGTACCGCTTGATAACGATTTGCTGAACACACCA<br>GTGTAAGGGATGTTTATGACGAGCAAAGAAACCTTTACCCATTACCAGCCGCAGGGCAACAGTGACCCGGCTCATACCGCAACCGCGCCCGGCGG<br>ATTGAGTGCGAAAGCGCCTGCAATGACCCCGCTGATGCTGGACACCTCCAGCCGTAAGCTGGTTGCGTGGGATGGCACCACCGACGGTGCTGCCG<br>TTGGCATTCTTGCGGTTGCTGCTCGAGCCTCAGCTCATGTCATCCTCAGCACACTTGACCCTCAGCTCAGCTAGCCTCAGCCTACAATCACCTCA<br>GCGAATTCGGTGACCCTTACGCGAATCCGCTTTCAGACGTTGACTGGTCGCGTCTGGCAAAAGTTAAAGACCTGACGCCCGGCGAACTGACCGCT<br>GAGTCCTATGACGACAGCTATCTCGATGATGAAGATGCAGACTGGACTGCGACCGGGCAGGGGCAGAAATCTGCCGGAGATACCAGCTTCACGCT<br>GGCGTGGATGCCCGGAGAGCAGGGGCAGCAGGCGCTGCTGGCGTGGTTTAATGAAGGCGATACCCGTGCCTATAAAATCCGCTTCCCGAACGGCA<br>CGGTCGATGTGTTCCGTGGCTGGGTCAGCAGTATCGGTAAGGCGGTGACGGCGAAGGAAGTGATCACCCGCACGGTGAAAGTCACCAATGTGGGA<br>CGTAACGATTTGCTGAACACACCAGTGTAAGGGATGTTTATGACGAGCAAAGAAACCTTTACCCATTACCAGCCGCAGGGCAACAGTGACCCGGC<br>TCATACCGCAACCGCGCCCGGCGGATTGAGTGCGAAAGCGCCTGCAATGACCCCGCTGATGCTGGACACCTCCAGCCGTAAGCTGGTTGCGTGGG<br>ATGGCACCACCGACGGTGCTGCCGTTGGCATTCTTGCGGTTGCTGCTCGAGCCTCAGCTCATGTCATCCTCAGCACACTTGACCCTCAGCTCAGC<br>TAGCCTCAGCCTACAATCACCTCAGCGAATTCGGTGACCCTTACGCGAATCCGCTTTCAGACGTTGACTGGTCGCGTCTGGCAAAAGTTAAAGAC<br>CTGACGCCCGGCGAACTGACCGCTGAGTCCTATGACGACAGCTATCTCGATGATGAAGATGCAGACTGGACTGCGACCGGGCAGGGGCAGAAATC<br>TGCCGGAGATACCAGCTTCACGCTGGCGTGGATGCCCGGAGAGCAGGGGCAGCAGGCGCTGCTGGCGTGGTTTAATGAAGGCGATACCCGTGCCT<br>ATAAAATCCGCTTCCCGAACGGCACGGTCGATGTGTTCCGTGGCTGGGTCAGCAGTATCGGTAAGGCGGTGACGGCGAAGGAAGTGATCACCCGC<br>ACGGTGAAAGTCACCAATGTGGGACGTATCGAATTCCTGCAGCCCGGGGGATCCACTAGTTCTAGAGCGGCCGCCACCGCGGTGGAGCTCCAGCT<br>TTTGTTCCCTTTAGTGAGGGTTAATTGCGCGCTTGGCGTAATCATGGTCATAGCTGTTTCCTGTGTGAAATTGTTATCCGCTCACAATTCCACAC<br>AACATACGAGCCGGAAGCATAAAGTGTAAAGCCTGGGGTGCCTAATGAGTGAGCTAACTCACATTAATTGCGTTGCGCTCACTGCCCGCTTTCCA<br>GTCGGGAAACCTGTCGTGCCAGCTGCATTAATGAATCGGCCAACGCGCGGGGAGAGGCGGTTTGCGTATTGGGCGCTCTTCCGCTTCCTCGCTCA<br>CTGACTCGCTGCGCTCGGTCGTTCGGCTGCGGCGAGCGGTATCAGCTCACTCAAAGGCGGTAATACGGTTATCCACAGAATCAGGGGATAACGCA<br>GGAAAGAACATGTGAGCAAAAGGCCAGCAAAAGGCCAGGAACCGTAAAAAGGCCGCGTTGCTGGCGTTTTTCCATAGGCTCCGCCCCCCTGACGA<br>GCATCACAAAAATCGACGCTCAAGTCAGAGGTGGCGAAACCCGACAGGACTATAAAGATACCAGGCGTTTCCCCCTGGAAGCTCCCTCGTGCGCT<br>CTCCTGTTCCGACCCTGCCGCTTACCGGATACCTGTCCGCCTTTCTCCCTTCGGGAAGCGTGGCGCTTTCTCATAGCTCACGCTGTAGGTATCTC<br>AGTTCGGTGTAGGTCGTTCGCTCCAAGCTGGGCTGTGTGCACGAACCCCCCGTTCAGCCCGACCGCTGCGCCTTATCCGGTAACTATCGTCTTGA<br>GTCCAACCCGGTAAGACACGACTTATCGCCACTGGCAGCAGCCACTGGTAACAGGATTAGCAGAGCGAGGTATGTAGGCGGTGCTACAGAGTTCT<br>TGAAGTGGTGGCCTAACTACGGCTACACTAGAAGGACAGTATTTGGTATCTGCGCTCTGCTGAAGCCAGTTACCTTCGGAAAAAGAGTTGGTAGC<br>TCTTGATCCGGCAAACAAACCACCGCTGGTAGCGGTGGTTTTTTTGTTTGCAAGCAGCAGATTACGCGCAGAAAAAAAGGATCTCAAGAAGATCC<br>TTTGATCTTTTCTACGG |
| C-trap central fragment<br><br>(25427 bp) | CGCGCATTGCGCCCAGCGCCATCTGATCGTTGGCAACCAGCATCGCAGTGGGAACGATGCCCTCATTCAGCATTTGCATGGTTTGTTGAAAACCG<br>GACATGGCACTCCAGTCGCCTTCCCGTTCCGCTATCGGCTGAATTTGATTGCGAGTGAGATATTTATGCCAGCCAGCCAGACGCAGACGCGCCGA<br>GACAGAACTTAATGGGCCCGCTAACAGCGCGATTTGCTGGTGACCCAATGCGACCAGATGCTCCACGCCCAGTCGCGTACCGTCTTCATGGGAGA<br>AAATAATACTGTTGATGGGTGTCTGGTCAGAGACATCAAGAAATAACGCCGGAACATTAGTGCAGGCAGCTTCCACAGCAATGGCATCCTGGTCA<br>TCCAGCGGATAGTTAATGATCAGCCCACTGACGCGTTGCGCGAGAAGATTGTGCACCGCCGCTTTACAGGCTTCGACGCCGCTTCGTTCTACCAT<br>CGACACCACCACGCTGGCACCCAGTTGATCGGCGCGAGATTTAATCGCCGCGACAATTTGCGACGGCGCGTGCAGGGCCAGACTGGAGGTGGCAA<br>CGCCAATCAGCAACGACTGTTTGCCCGCCAGTTGTTGTGCCACGCGGTTGGGAATGTAATTCAGCTCCGCCATCGCCGCTTCCACTTTTTCCCGC<br>GTTTTCGCAGAAACGTGGCTGGCCTGGTTCACCACGCGGGAAACGGTCTGATAAGAGACACCGGCATACTCTGCGACATCGTATAACGTTACTGG<br>TTTCACATTCACCACCCTGAATTGACTCTCTTCCGGGCGCTATCATGCCATACCGCGAAAGGTTTTGCGCCATTCGATGGTGTCCGGGATCTCGA<br>CGCTCTCCCTTATGCGACTCCTGCATTAGGAAGCAGCCCAGTAGTAGGTTGAGGCCGTTGAGCACCGCCGCCGCAAGGAATGGTGCATGCAAGGA<br>GATGGCGCCCAACAGTCCCCCGGCCACGGGGCCTGCCACCATACCCACGCCGAAACAAGCGCTCATGAGCCCGAAGTGGCGAGCCCGATCTTCCC<br>CATCGGTGATGTCGGCGATATAGGCGCCAGCAACCGCACCTGTGGCGCCGGTGATGCCGGCCACGATGCGTCCGGCGTAGAGGATCGAGATCTCG<br>ATCCCGCGAAATTAATACGACTCACTATAGGGGAATTGTGAGCGGATAACAATTCCCTTCTAGCCATATTGGACTCGGACCTGTTTCACGTGGAA<br>CACTGAGCCTGGACTAGGTCTAGCATATTGGACTCGACTTTCACGTGGAACACTGAGCCTGGACTAGGTCTAGAACAGGTTCTTTTTTCTTTGTT<br>TCACGTGGAACATTCTGATTAATGTACAGCTAGCCATATTGGACTCGGACCTGTTTCACGTGGAACACTGAGCCTGGACTAGGTCTAGAACAGGT<br>TCTTTTTTCTTTGTTTCACGTGGAACATTCTGATTAATGTACAGCTAGAACAGGTTCTTTTTTCTTTGTTTCACGTGGAACATTCTGATTAATGT<br>ACAGCTAGAAATAATTTTGTTTAACTTTAAGAAGGAGATATACCATGTGTTTCACGTGGAACACATGGCTAAAGGCCTTGGAAAAGGGATTAATG<br>CGTTATTTAATCAGGTAGATTTGTCTGAAGAGACAGTTGAAGAAATTAAAATTGCCGATTTACGCCCTAATCCTTATCAGCCAAGAAAACACTTT<br>GATGACGAGGCATTAGCTGAACTAAAAGAATCTGTGCTGCAGCATGGCATTCTTCAGCCGCTTATCGTCAGAAAATCTTTAAAAGGCTATGATAT<br>TGTTGCGGGTGAACGGCGTTTTCGAGCGGCAAAGCTGGCAGGTTTAGATACAGTTCCGGCCATTGTCCGTGAATTATCAGAGGCGTTAATGAGGG<br>AAATTGCTTTATTAGAAAACCTTCAGCGTGAAGATTTATCTCCGCTTGAAGAGGCTCAGGCATATGCCGAAACAAGCGCTCATGAGCCCGAAGTG<br>GCGAGCCCGATCTTCCCCATCGGTGATGTCGGCGATATAGGCGCCAGCAACCGCACCTGTGGCGCCGGTGATGCCGGCCACGATGCGTCCGGCGT<br>AGAGGATCGAGATCTCGATCCCGCGAAATTAATACGACTCACTATAGGGGAATTGTGAGCGGATAACAATTCCCTTCTAGCCATATTGGACTCGG<br>ACCTGTTTCACGTGGAACACTGAGCCTGGACTAGGTCTAGCATATTGGACTCGACTTTCACGTGGAACACTGAGCCTGGACTAGGTCTAGAACAG<br>GTTCTTTTTTCTTTGTTTCACGTGGAACATTCTGATTAATGTACAGCTAGCCATATTGGACTCGGACCTGTTTCACGTGGAACACTGAGCCTGGA<br>CTAGGTCTAGAACAGGTTCTTTTTTCTTTGTTTCACGTGGAACATTCTGATTAATGTACAGCTAGAACAGGTTCTTTTTTCTTTGTTTCACGTGG<br>AACATTCTGATTAATGTACAGCTAGAAATAATTTTGTTTAACTTTAAGAAGGAGATATACCATGTGTTTCACGTGGAACACATGGCTAAAGGCCT<br>TGGAAAAGGGATTAATGCGCATATGACTCCCTTTTGAAACACTTAGATCTCACACAAGAGCAGCTTGCCAAACGTCTTGGGAAAAGCAGACCGCA<br>TATTGCGAATCATTTAAGACTGCTGACACTGCCAGAAAATATTCAACAGCTTATTGCCGAAGGCACGCTTTCTATGGGACATGGACGCACGCTTC |

| | |
|---|---|
| | TTGGCTTAAAAAACAAAAATAAGCTTGAACCGCTGGTACAAAAAGTGATTGCGGAGCAGCTCAATGTTCGCCAACTTGAGCAGCTGATTCAGCAG<br>TTGAATCAGAATGTTCCACGTGAAACAAAGAAAAAAGAACCTGTGAAAGATGCGGTTCTAAAAGAACGGGAATCCTATCTCCAAAATTATTTGG<br>AACAACAGTTAATATTAAAAGACAGAAGAAAAAAGGCAAAATCGAAATTGAATTTTTCTCTAATGAAGACCTTGACCGGATTTTAGAGCTTTTGT<br>CTGAACGAGAATCATAAGGATCCGAATTCATCCGAGAATTCCGATAGGAATTCCACGTTGAATTCTACGACGAATTCTCAACTGTGAGGAGGCCA<br>CGGTACTGGAGTCGTTTCTGGAAGAGCACGGGGGCTGGAAATCCTTTCTGTGGACGCCGCCTTATGAGTGGCGGCAGATAAAGGTGACCTGCGCA<br>AAATGGTCGTCGCGGGTCAGTATGCTGCGTGTTGAGTTCAGCGCAGAGTTTGAACAGGTGGTGAACTGATGCAGGATATCCGGCAGGAAACACTG<br>AATGAATGCACCCGTGCGGAGCAGTCGGCCAGCGTGGTGCTCTGGGAAATCGACCTGACAGAGGTCGGTGGAGAACGTTATTTTTTCTGTAATGA<br>GCAGAACGAAAAAGGTGAGCCGGTCACCTGGCAGGGGCGACAGTATCAGCCGTATCCCATTCAGGGGAGCGGTTTTGAACTGAATGGCAAAGGCA<br>CCAGTACGCGCCCCACGCTGACGGTTTCTAACCTGTACGGTATGGTCACCGGGATGGCGGAAGATATGCAGAGTCTGGTCGGCGGAACGGTGGTC<br>CGGCGTAAGGTTTACGCCCGTTTTCTGGATGCGGTGAACTTCGTCAACGGAAACAGTTACGCCGATCCGGAGCAGGAGGTGATCAGCCGCTGGCG<br>CATTGAGCAGTGCAGCGAACTGAGCGCGGTGAGTGCCTCCTTTGTACTGTCCACGCCGACGGAAACGGATGGCGCTGTTTTTCCGGGACGTATCA<br>TGCTGGCCAACACCTGCACCTGGACCTATCGCGGTGACGAGTGCGGTTATAGCGGTCCGGCTGTCGCGGATGAATATGACCAGCCAACGTCCGAT<br>ATCACGAAGGATAAATGCAGCAAATGCCTGAGCGGTTGTAAGTTCCGCAATAACGTCGGCAACTTTGGCGGCTTCCTTTCCATTAACAAACTTTC<br>GCAGTAAATCCCATGACACAGACAGAATCAGCGATTCTGGCGCACGCCCGGCGATGTGCGCCAGCGGAGTCGTGCGGCTTCGTGGTAAGCACGCC<br>GGAGGGGGAAAGATATTTCCCCTGCGTGAATATCTCCGGTGAGCCGGAGGCTATTTCCGTATGTCGCCGGAAGACTGGCTGCAGGCAGAAATGCA<br>GGGTGAGATTGTGGCGCTGGTCCACAGCCACCCCGGTGGTCTGCCCTGGCTGAGTGAGGCCGACCGGCGGCTGCAGGTGCAGAGTGATTTGCCGT<br>GGTGGCTGGTCTGCCGGGGGACGATTCATAAGTTCCGCTGTGTGCCGCATCTCACCGGGCGGCGCTTTGAGCACGGTGTGACGGACTGTTACACA<br>CTGTTCCGGGCGGCCCCGCAAGGATTGCCCCGATGCCTTGTTCCTTTGCCGCGAGAATGGCGGCCAACAGGTCATGTTTTTCTGGCATCTTCATG<br>TCTTACCCCCAATAAGGGGATTTGCTCTATTTAATTAGGAATAAGGTCGATTACTGATAGAACAAATCCAGGCTACTGTGTTTAGTAATCAGATT<br>TGTTCGTGACCGATATGCACGGGCAAAACGGCAGGAGGTTGTTAGCGCGACCTCCTGCCACCCGCTTTCACGAAGGTCATGTGTAAAAGGCCGCA<br>GCGTAACTATTACTAATGAATTCAGGACAGACAGTGGCTACGGCTCAGTTTGGGTTGTGCTGTTGCTGGGCGGCGATGACGCCTGTACGCATTTG<br>GTGATCCGGTTCTGCTTCCGGTATTCGCTTAATTCAGCACAACGGAAAGAGCACTGGCTAACCAGGCTCGCCGACTCTTCACGATTATCGACTCA<br>ATGCTCTTACCTGTTGTGCAGATATAAAAAATCCCGAAACCGTTATGCAGGCTCTAACTATTACCTGCGAACTGTTTCGGGATTGCATTTTGCAG<br>ACCTCTCTGCCTGCGATGGTTGGAGTTCCAGACGATACGTCGAAGTGACCAACTAGGCGGAATCGGTAGTAAGCGCCGCCTCTTTTCATCTCACT<br>ACCACAACGAGCGAATTAACCCATCGTTGAGTCAAATTTACCCAATTTTATTCAATAAGTCAATATCATGCCGTTAATATGTTGCCATCCGTGGC<br>AATCATGCTGCTAACGTGTGACCGCATTCAAAATGTTGTCTGCGATTGACTCTTCTTTGTGGCATTGCACCACCAGAGCGTCATACAGCGGCTTA<br>ACAGTGCGTGACCAGGTGGGTTGGGTAAGGTTTGGGATTAGCATCGTCACAGCGCGATATGCTGCGCTTGCTGGCATCCTTGAATAGCCGACGCC<br>TTTGCATCTTCCGCACTCTTTCTCGACAACTCTCCCCCACAGCTCTGTTTTGGCAATATCAACCGCACGGCCTGTACCATGGCAATCTCTGCATC<br>TTGCCCCCGGCGTCGCGGCACTACGGCAATAATCCGCATAAGCGAATGTTGCGAGCACTTGCAGTACCTTTGCCTTAGTATTTCCTTCAAGCTTT<br>GCCACACCACGGTATTTCCCCGATACCTTGTGTGCAAATTGCATCAGATAGTTGATAGCCTTTTGTTTGTCGTTCTGGCTGAGTTCGTGCTTACC<br>GCAGAATGCAGCCATACCGAATCCGGCTTGTGATTGCGCCATCCCCATAGCAGCCATCACATCAGTACCGGAAAGAGAGTCAGAAGCCGTGGCCC<br>GTGGTGAGTCGCTCATCATCGGGCTTTTTGGCGAATGAAATTTAGCTACGCTTTCGAGTCTCATGCGCCTTCTCCCTGTACCTGAATCAATGTTA<br>GGTTTCCGCAGAACACTGCGCCGGTATCGATATACATTTGGTTGGCAAACTTGAGTGGTTTCACTGCTGGCGTATGACCAAAGATGAACGTGTCC<br>GCGCCTTTGATTTCTTTCACGATCCCGTTTTGTGAGTTGCTGATTCGTTCGCGGTTCCAGATTACCTGCTGATGATCAACTGGCTTTCCAAACTC<br>GTATTCGTCAAAGGGATAATCGGCGTGGCAGATAACATATTTTTTATCTTTGCTCACCAGTTCGATGATTAACGGAAGTTCATCTGCTTTATGGG<br>CAAGAGCTTTAGCCAGAATTTCTTTGTCGTAATCGAGATTAAAGAACCAGCCACCGCCATTAAGCAGCCAGTGATTAACGTTTCCACGCTCTGAT<br>AAGCCATCAATCATCATTTGCTCATGGTTTCCACGTACAGCTCTGAACCAGGGGAATGTGATTAATTCCAGGCATTCAACGTTCTCTGCACCACG<br>ATCAACCAAATCGCCCACCGAGATAAGCAGGTCTTTTTTGTTGTCGAATCCAATCGTATCCAGTTTGTTCATCAGGTTCGTGTAGCATCCGTGCA<br>GATCGCCAACTACCCAAATATTTCGGTATTTGCTGCCATCAATTTTTTCGTAATAGCGCATCTCTTTCACTCCATCCGCGATGAACCATGAGAAC<br>GTCGTTGACGATGGCGTGCATTTTCCCGTCTTTATCATCAACGTATTTTCTGACCGTACCGCGACTACATTTCAGTCTGCGTGCTACTTCTGTCT<br>GATTTCCGTATGTTTCAACGAGCATGTCTGGAATGGTTTTTACTGAGAACGTCATGCGGCCTCACTTCTGCTATTTCGCAGGTCTTTGAGTTTCT<br>GTTGGTACTCTGCCTTGATCGCCTTGCACTCTTCGATAGTCCAGCGATGGCGGTTATGGTTTGATTCGATTTCGTCTACTGCTTCCTGCCCGATG<br>CGGCTAATCAGTTCGACGCGATACGGAACGAGATTTCCGCTTTTGTGCTGGTTGCACACCACGCATTGCTTGTGAATATTGCGTTCATTAAATCG<br>GAGTTGAGGTGCCGCAGCAGTTGTCCGGTAATGTCCGGCATCCCACTGAGCAGACGTGAGCGTTCCGCACGAGATACATGGTAAGTCGCGGTCTC<br>TTTCTCTGATGAAGGCGTTTACGGCTTGTTGGGCTTGTTTAATCCAGTAACTGCGGGGCTTTAAGGCGAGTTTTCGAATCTTAAGTTTATCTTTC<br>TGTTTCTGCTCCTCTCGTCGTCGTTTCTTCTCTGCTGCTTTTTCCGCTTTTTCGCGTTCTTTACTTCGTCGTTCGAGTGCTATCTTGGTTCCACA<br>CTCTGGAGAGCACCACCACTGATTAGCGAATGCAGGGTGAAACCATTCCCGGCATTCATCGTTTTTACATCGTCTTCGCGCTGGTTTAGCCATCA<br>TCTTCTTCCTCGTGCATCGAGCTATTCGGATCGCTCATCAGTTCTGCGCAGCAGTGCTCACACACGTGAACTTCCAGCACATGCAGCTTCTGACC<br>GCAGTTAGCGCACGTTAAAGCTCGCTCGACGCTTTCTTGTTCGTAACTTCGATTTTGGTCAATCACCTTGTTTTCCTCGCACGACGTCTTAGCCA<br>CCGGATATCCCACAGGTGAGCCGTGTAGTTGAAGGTTTTTACGTCAGATTCTTTTGGGATTGGCTTGGGTTTATTTCTGGTGCGTTTCGTTGGAA<br>GGTATTTGCAGTTTTCGCAGATTATGTCGGTGATACTTCGTCGCTGTCTCGCCACACGTCCTCCTTTTCCTGCGGTAGTGGTAACACCCCTGTTG<br>GTGTTCTTTCACACCGGAGACACCATCGATTCCAGTAAGGTTGATTTGGTCGGAAGCGGTTATCTTCTTTGCATTCACCGCACCGATAACATCGC<br>ATCATGCAGCTTCCCTCCCGAAGTCGAAATCAAGCTGCCCTCCAAATATTTCGCATGACTCAGAACAAGAGCCGGTATCGAATCTTTTAGCTCGT<br>ACCATGTCCTGATACAGGGCTTGATAATCATTTTCTGAATACATTTTCGCGATACCGTCCAGCGACATTCTTCCTCGGTACATAATCTCCTTTGG<br>CGTTTCCCGATGTCCGTCACGCACATGGGATCCCGTGATGACCTCATTAAAAACACGCTGCAATCCCTCCTCATCTTTGCAGGCAAGTCCGATTT<br>TTTGCGTTGATTTTTTAATGCAGAATATGCAGTTACCGAGATGTTCCGGTATTTGCAAATCGAATGGTTGTTGCTTCCACCATGCGAGGATATCT<br>TCCTTCTCAAAGTCTGACAGTTCAGCAAGATATCTGATTCCAGGCTTTGGCTTTAGCCGCTTCGGTTCATCAGCTCTGATGCCAATCCACGTGGT<br>GTAATTCCCTCGCCCGAAATGGTCATCACAGTATTTGGTGAAGGGAACGAGTTTTAATCTGTCAGTGCAGAACGCGCCGCCGACGTATGGAGTGC<br>CATATTTCTTTACCATATCGATAAATGGCTTCAGAACAGGCATTCGCGTCTGAATATCCTTTGGTTCCCATACCGTATAACCATTTGGCTGTCCA<br>AGCTCCGGGTTGATATCAACCTGCAATACGGTGAGCGGTATATCCCAGAACTTCACAACTTCCCTGACAAACCGATATGTCATTGGATGTTCACA<br>ACCTGTATCCATGAAAACGTAATGCACGTCTTTACCTGCCCGTCGCTTTTGCTCCATTAGCCAGAGCAAATATGCTGACGTCCTGCCACCGGAGA<br>AACTAACGACATTTATCATGCAGCCCTGTCTCCCCATCTCGCTTTCCACTCCAGAGCCAGTCTCGCTTCGTCTGACCACTTAACGCCACGCTCTG<br>TACCGAATGCCTGTATAAGCTCTAATAGCTCCGCAAATTCGCCTACACGCATCCTGCTGGTTGACTGGCCTATTACCACAAAGCCATTCCCGGCA<br>AGGTTAGGAACAACATCCTGCTGCTTTAATGCTGCGGTAAACACACACTTCCAGCTTTCTGCATCCAGCCAGCGACCATGCCATTCAACCTGACG<br>AGAGACGTCACCTAAGCAGGCCCATAGCTTCCTGTTTTGGTCTAAGCTGCGGTTGCGTTCCTGAATGGTTACTACGATTGGTTTGGTTGGGTCTG<br>GAAGGATTTGCTGTACTGCGTGAATAGCGTTTTGCTGATGTGCTGGAGATCGAATTTCAAAGGTTAGTTTTTTCATGACTTCCCTCTCCCCCAAA<br>TAAAAAGGCCTGCGATTACCAGCAGGCCTGTTATTAGCTCAGTAATGTAGATGGTCATCTTTTAACTCCATATACCGCCAATACCCGTTTCATCG<br>CGGCACTCTGGCGACACTCCTTAAAAACCAGGTTCGTGCTCATCTTTCCTTCCCGTTCTTCCCTGGTAGCAAACCGGTAATACACCGTTCGCCAG<br>ACCTTACCTTCGATAACCAGAAGACCTGCCCGTGCCATTTTAGCCGCGGCCTGATTTATGCTGGTTACTGTTGCGCCTGTTAGCGCGGCAACGTC<br>CGGCGCACAGAAGCTATTATGCGTCCCCAGGTAATGAATAATTGCCTCTTTGCCCGTCATACACTTGCTCCTTTCAGTCCGAACTTAGCTTTGAT<br>TTCTGCGATCTTCGCCAGAGCCTGTGCACGATTTAGAGGTCTACCGCCCATGACAGGAAGTTGTTTTACTGGTTCAGGGATCGCCTCACCACGGT<br>TAATTCTCGCAGTCATATGGACAAGCTCATCTGCGGCCTTACGGCGTAATTCCGCATCAGTAAGCGCATTGGCCCGCATGTTCTGATACAGGTTG<br>GTAACCAGCCAGTAGTGCGCGTTTGATTTCCACGGATAAGACTCCGCATCCGGCTCGAGCCTCAGCTCATGTCATCCTCAGCACACTTGACCCTC<br>AGCTCAGCTAGCCTCAGCCTACAATCACCTCAGCGAATTCGGTGACCCTTACGCGAATCCGCTTTCAGACGTTGACTGGTCGCGTCTGGCAAAAG<br>TTAAAGACCTGACGCCCGGCGAACTGACCGCTGAGTCCTATGACGACAGCTATCTCGATGATGAAGATGCAGACTGGACTGCGACCGGGCAGGGG<br>CAGAAATCTGCCGGAGATACCAGCTTCACGCTGGCGTGGATGCCCGGAGAGCAGGGGCAGCAGGCGCTGCTGGCGTGGTTTAATGAAGGCGATAC<br>CCGTGCCTATAAAATCCGCTTCCCGAACGGCACGGTCGATGTGTTCCGTGGCTGGGTCAGCAGTATCGGTAAGGCGGTGACGGCGAAGGAAGTGA<br>TCACCCGCACGGTGAAAGTCACCAATGTGGGACGTCCGTCGATGGCAGAAGATCGCAGCACGGTAACAGCGGCAACCGGCATGACCGTGACGCCT<br>GCCAGCACCTCGGTGGTGAAAGGGCAGAGCACCACGCTGACCGTGGCCTTCCAGCCGGAGGGCGTAACCGACAAGAGCTTTCGTGCGGTGTCTGC<br>GGATAAAACAAAAGCCACCGTGTCGGTCAGTGGTATGACCATCACCGTGAACGGCGTTGCTGCAGGCAAGGTCAACATTCCGGTTGTATCCGGTA<br>ATGGTGAGTTTGCTGCGGTTGCAGAAATTACCGTCACCGCCAGTTAATCCGGAGAGTCAGCGATGTTCCTGAAAACCGAATCATTTGAACATAAC<br>GGTGTGACCGTCACGCTTTCTGAACTGTCAGCCCTGCAGCGCATTGAGCATCTCGCCCTGATGAAACGGCAGGCAGAACAGGCGGAGTCAGACAG<br>CAACCGGAAGTTTACTGTGGAAGACGCCATCAGAACCGGCGCGTTTCTGGTGGCGATGTCCCTGTGGCATAACCATCCGCAGAAGACGCAGATGC<br>CGTCCATGAATGAAGCCGTTAAACAGATTGAGCAGGAAGTGCTTACCACCTGGCCCACGGAGGCAATTTCTCATGCTGAAAACGTGGTGTACCGG<br>CTGTCTGGTATGTATGAGTTTGTGGTGAATAATGCCCCTGAACAGACAGAGGACGCCGGGCCCGCAGAGCCTGTTTCTGCGGGAAAGTGTTCGAC<br>GGTGAGCTGAGTTTTGCCCTGAAACTGGCGCGTGAGATGGGGCGACCCGACTGGCGTGCCATGCTTGCCGGGATGTCATCCACGGAGTATGCCGA<br>CTGGCACCGCTTTTACAGTACCCATTATTTTCATGATGTTCTGCTGGATATGCACTTTTCCGGGCTGACGTACACCGTGCTCAGCCTGTTTTTCA |

GCGATCCGGATATGCATCCGCTGGATTTCAGTCTGCTGAACCGGCGCGAGGCTGACGTCGAGTGAAGCGTTATTGGTATGCGGTAAAACCGCACT
CAGGCGGCCTTGATAGTCATATCATCTGAATCAAATATTCCTGATGTATCGATATCGGTAATTCTTATTCCTTCGCTACCATCCATTGGAGGCCA
TCCTTCCTGACCATTTCCATCATTCCAGTCGAACTCACACACAACACCATATGCATTTAAGTCGCTTGAAATTGCTATAAGCAGAGCATGTTGCG
CCAGCATGATTAATACAGCATTTAATACAGAGCCGTGTTTATTGAGTCGGTATTCAGAGTCTGACCAGAAATTATTAATCTGGTGAAGTTTTTCC
TCTGTCATTACGTCATGGTCGATTTCAATTTCTATTGATGCTTTCCAGTCGTAATCAATGATGTATTTTTTGATGTTTGACATCTGTTCATATCC
TCACAGATAAAAAATCGCCCTCACACTGGAGGGCAAAGAAGATTTCCAATAATCAGAACAAGTCGGCTCCTGTTTAGTTACGAGCGACATTGCTC
CGTGTATTCACTCGTTGGAATGAATACACAGTGCAGTGTTTATTCTGTTATTTATGCCAAAAATAAAGGCCACTATCAGGCAGCTTTGTTGTTCT
GTTTACCAAGTTCTCTGGCAATCATTGCCGTCGTTCGTATTGCCCATTTATCGACATATTTCCCATCTTCCATTACAGGAAACATTTCTTCAGGC
TTAACCATGCATTCCGATTGCAGCTTGCATCCATTGCATCGCTTGAATTGTCCACACCATTGATTTTTATCAATAGTCGTAGTCATACGGATAGT
CCTGGTATTGTTCCATCACATCCTGAGGATGCTCTTCGAACTCTTCAAATTCTTCTTCCATATATCACCTTAAATAGTGGATTGCGGTAGTAAAG
ATTGTGCCTGTCTTTTAACCACATCAGGCTCGGTGGTTCTCGTGTACCCCTACAGCGAGAAATCGGATAAACTATTACAACCCCTACAGTTTGAT
GAGTATAGAAATGGATCCACTCGTTATTCTCGGACGAGTGTTCAGTAATGAACCTCTGGAGAGAACCATGTATATGATCGTTATCTGGGTTGGAC
TTCTGCTTTTAAGCCCAGATAACTGGCCTGAATATGTTAATGAGAGAATCGGTATTCCTCATGTGTGGCATGTTTTCGTCTTTGCTCTTGCATTT
TCGCTAGCAATTAATGTGCATCGATTATCAGCTATTGCCAGCGCCAGATATAAGCGATTTAAGCTAAGAAAACGCATTAAGATGCAAAACGATAA
AGTGCGATCAGTAATTCAAAACCTTACAGAAGAGCAATCTATGGTTTTGTGCGCAGCCCTTAATGAAGGCAGGAAGTATGTGGTTACATCAAAAC
AATTCCCATACATTAGTGAGTTGATTGAGCTTGGTGTGTTGAACAAAACTTTTTCCCGATGGAATGGAAAGCATATATTATTCCCTATTGAGGAT
ATTTACTGGACTGAATTAGTTGCCAGCTATGATCCATATAATATTGAGATAAAGCCAAGGCCAATATCTAAGTAACTAGATAAGAGGAATCGATT
TTCCCTTAATTTTCTGGCGTCCACTGCATGTTATGCCGCGTTCGCCAGGCTTGCTGTACCATGTGCGCTGATTCTTGCGCTCAATACGTTGCAGG
TTGCTTTCAATCTGTTTGTGGTATTCAGCCAGCACTGTAAGGTCTATCGGATTTAGTGCGCTTTCTACTCGTGATTTCGGTTTGCGATTCAGCGA
GAGAATAGGGCGGTTAACTGGTTTTGCGCTTACCCCAACCAACAGGGGATTTGCTGCTTTCCATTGAGCCTGTTTCTCTGCGCGACGTTCGCGGC
GGCGTGTTTGTGCATCCATCTGGATTCTCCTGTCAGTTAGCTTTGGTGGTGTGTGGCAGTTGTAGTCCTGAACGAAAACCCCCCGCGATTGGCAC
ATTGGCAGCTAATCCGGAATCGCACTTACGGCCAATGCTTCGTTTCGTATCACACACCCCAAAGCCTTCTGCTTTGAATGCTGCCCTTCTTCAGG
GCTTAATTTTTAAGAGCGTCACCTTCATGGTGGTCAGTGCGTCCTGCTGATGTGCTCAGTATCACCGCCAGTGGTATTTATGTCAACACCGCCAG
AGATAATTTATCACCGCAGATGGTTATCTGTATGTTTTTTATATGAATTTATTTTTTGCAGGGGGGCATTGTTTGGTAGGTGAGAGATCTGAATT
GCTATGTTTAGTGAGTTGTATCTATTTATTTTTCAATAAATACAATTGGTTATGTGTTTTGGGGGCGATCGTGAGGCAAAGAAAACCCGGCGCTG
AGGCCGGGTTATTCTTGTTCTCTGGTCAAATTATATAGTTGGAAAACAAGGATGCATATATGAATGAACGATGCAGAGGCAATGCCGATGGCGAT
AGTGGGTATCATGTAGCCGCTTATGCTGGAAAGAAGCAATAACCCGCAGAAAAACAAAGCTCCAAGCTCAACAAAACTAAGGGCATAGACAATAA
CTACCGATGTCATATACCCATACTCTCTAATCTTGGCCAGTCGGCGCGTTCTGCTTCCGATTAGAAACGTCAAGGCAGCAATCAGGATTGCAATC
ATGGTTCCTGCATATGATGACAATGTCGCCCCAAGACCATCTCTATGAGCTGAAAAAGAAACACCAGGAATGTAGTGGCGGAAAAGGAGATAGCA
AATGCTTACGATAACGTAAGGAATTATTACTATGTAAACACCAGGCATGATTCTGTTCCGCATAATTACTCCTGATAATTAATCCTTAACTTTGC
CCACCTGCCTTTTAAAACATTCCAGTATATCACTTTTCATTCTTGCGTAGCAATATGCCATCTCTTCAGCTATCTCAGCATTGGTGACCTTGTTC
AGAGGCGCTGAGAGATGGCCTTTTTCTGATAGATAATGTTCTGTTAAAATATCTCCGGCCTCATCTTTTGCCCGCAGGCTAATGTCTGAAAATTG
AGGTGACGGGTTAAAAATAATATCCTTGGCAACCTTTTTTATATCCCTTTTAAATTTTGGCTTAATGACTATATCCAATGAGTCAAAAAGCTCCC
CTTCAATATCTGTTGCCCCTAAGACCTTTAATATATCGCCAAATACAGGTAGCTTGGCTTCTACCTTCACCGTTGTTCGGCCGATGAAATGCATA
TGCATAACATCGTCTTTGGTGGTTCCCCTCATCAGTGGCTCTATCTGAACGCGCTCTCCACTGCTTAATGACATTCCTTTCCCGATTAAAAAATC
TGTCAGATCGGATGTGGTCGGCCCGAAAACAGTTCTGGCAAAACCAATGGTGTCGCCTTCAACAAACAAAAAAGATGGGAATCCCAATGATTCGT
CATCTGCGAGGCTGTTCTTAATATCTTCAACTGAAGCTTTAGAGCGATTTATCTTCTGAACCAGACTCTTGTCATTTGTTTTGGTAAAGAGAAAA
GTTTTTCCATCGATTTTATGAATATACAAATAATTGGAGCCAACCTGCAGGTGATGATTATCAGCCAGCAGAGAATTAAGGAAAACAGACAGGTT
TATTGAGCGCTTATCTTTCCCTTTATTTTTGCTGCGGTAAGTCGCATAAAAACCATTCTTCATAATTCAATCCATTTACTATGTTATGTTCTGAG
GGGAGTGAAAATTCCCCTAATTCGATGAAGATTCTTGCTCAATTGTTATCAGCTATGCGCCGACCAGAACACCTTGCCGATCAGCCAAACGTCTC
TTCAGGCCACTGACTAGCGATAACTTTCCCCACAACGGAACAACTCTCATTGCATGGGATCATTGGGTACTGTGGGTTTAGTGGTTGTAAAAACA
CCTGACCGCTATCCCTGATCAGTTTCTTGAAGGTAAACTCATCACCCCCAAGTCTGGCTATGCAGAAATCACCTGGCTCAACAGCCTGCTCAGGG
TCAACGAGAATTAACATTCCGTCAGGAAAGCTTGGCTTGGAGCCTGTTGGTGCGGTCATGGAATTACCTTCAACCTCAAGCCAGAATGCAGAATC
ACTGGCTTTTTTGGTTGTGCTTACCCATCTCTCCGCATCACCTTTGGTAAAGGTTCTAAGCTTAGGTGAGAACATCCCTGCCTGAACATGAGAAA
AAACAGGGTACTCATACTCACTTCTAAGTGACGGCTGCATACTAACCGCTTCATACATCTCGTAGATTTCTCTGGCGATTGAAGGGCTAAATTCT
TCAACGCTAACTTTGAGAATTTTTGTAAGCAATGCGGCGTTATAAGCATTTAATGCATTGATGCCATTAAATAAAGCACCAACGCCTGACTGCCC
CATCCCCATCTTGTCTGCGACAGATTCCTGGGATAAGCCAAGTTCATTTTTCTTTTTTTCATAAATTGCTTTAAGGCGACGTGCGTCCTCAAGCT
GCTCTTGTGTTAATGGTTTCTTTTTTGTGCTCATACGTTAAATCTATCACCGCAAGGGATAAATATCTAACACCGTGCGTGTTGACTATTTTACC
TCTGGCGGTGATAATGGTTGCATGTACTAAGGAGGTTGTATGGAACAACGCATAACCCTGAAAGATTATGCAATGCGCTTTGGGCAAACCAAGAC
AGCTAAAGATCTCGGCGTATATCAAAGCGCGATCAACAAGGCCATTCATGCAGGCCGAAAGATTTTTTTAACTATAAACGCTGATGGAAGCGTTT
ATGCGGAAGAGGTAAAGCCCTTCCCGAGTAACAAAAAAACAACAGCATAAATAACCCCGCTCTTACACATTCCAGCCCTGAAAAAGGGCATCAAA
TTAAACCACACCTATGGTGTATGCATTTATTTGCATACATTCAATCAATTGTTATCTAAGGAAATACTTACATATGGTTCGTGCAAACAAACGCA
ACGAGGCTCTACGAATCGAGAGTGCGTTGCTTAACAAAATCGCAATGCTTGGAACTGAGAAGACAGCGGAAGCTGTGGGCGTTGATAAGTCGCGA
ATTCGTCGTAGAATTCAACGTGGAATTCCTATCGGAATTCTCGGATGAATTCGTCGAGCCTCAGCTCATGTCATCCTCAGCACACTTGACCCTCA
GCTCAGCTAGCCTCAGCCTACAATCACCTCAGCGAATTCGGTGACCCTTACGCGAATCCGCTTTCAGACGTTGACTGGTCGCGTCTGGCAAAAGT
TAAAGACCTGACGCCCGGCGAACTGACCGCTGAGTCCTATGACGACAGCTATCTCGATGATGAAGATGCAGACTGGACTGCGACCGGGCAGGGGC
AGAAATCTGCCGGAGATACCAGCTTCACGCTGGCGTGGATGCCCGGAGAGCAGGGGCAGCAGGCGCTGCTGGCGTGGTTTAATGAAGGCGATACC
CGTGCCTATAAAATCCGCTTCCCGAACGGCACGGTCGATGTGTTCCGTGGCTGGGTCAGCAGTATCGGTAAGGCGGTGACGGCGAAGGAAGTGAT
CACCCGCACGGTGAAAGTCACCAATGTGGGACGTCCGTCGATGGCAGAAGATCGCAGCACGGTAACAGCGGCAACCGGCATGACCGTGACGCCTG
CCAGCACCTCGGTGGTGAAAGGGCAGAGCACCACGCTGACCGTGGCCTTCCAGCCGGAGGGCGTAACCGACAAGAGCTTTCGTGCGGTGTCTGCG
GATAAAACAAAAGCCACCGTGTCGGTCAGTGGTATGACCATCACCGTGAACGGCGTTGCTGCAGGCAAGGTCAACATTCCGGTTGTATCCGGTAA
TGGTGAGTTTGCTGCGGTTGCAGAAATTACCGTCACCGCCAGTTAATCCGGAGAGTCAGCGATGTTCCTGAAAACCGAATCATTTGAACATAACG
GTGTGACCGTCACGCTTTCTGAACTGTCAGCCCTGCAGCGCATTGAGCATCTCGCCCTGATGAAACGGCAGGCAGAACAGGCGGAGTCAGACAGC
AACCGGAAGTTTACTGTGGAAGACGCCATCAGAACCGGCGCGTTTCTGGTGGCGATGTCCCTGTGGCATAACCATCCGCAGAAGACGCAGATGCC
GTCCATGAATGAAGCCGTTAAACAGATTGAGCAGGAAGTGCTTACCACCTGGCCCACGGAGGCAATTTCTCATGCTGAAAACGTGGTGTACCGGC
TGTCTGGTATGTATGAGTTTGTGGTGAATAATGCCCCTGAACAGACAGAGGACGCCGGGCCCGCAGAGCCTGTTTCTGCGGAAAGTGTTCGACG
GTGAGCTGAGTTTTGCCCTGAAACTGGCGCGTGAGATGGGGCGACCCGACTGGCGTGCCATGCTTGCCGGGATGTCATCCACGGAGTATGCCGAC
TGGCACCGCTTTTACAGTACCCATTATTTTCATGATGTTCTGCTGGATATGCACTTTTCCGGGCTGACGTACACCGTGCTCAGCCTGTTTTTCAG
CGATCCGGATATGCATCCGCTGGATTTCAGTCTGCTGAACCGGCGCGAGGCTGACGTCGACAAGCTTCCGGCCGCAGTCGAGCACCACCACCACC
ACCACTGAGATCCGGCTGCTAACAAAGCCCGAAAGGAAGCTGAGTTGGCTGCTGCCACCGCTGAGCAATAACTAGCATAACCCCTTGGGGCCTCT
AAACGGGTCTTGAGGGGTTTTTTGCTGAAAGGAGGAACTATATCCGGATTGGCGAATGGGACGCGCCCTGTAGCGGCGCATTAAGCGCGGCGGGT
GTGGTGGTTACGCGCAGCGTGACCGCTACACTTGCCAGCGCCCTAGCGCCCGCTCCTTTCGCTTTCTTCCCTTCCTTTCTCGCCACGTTCGCCGG
CTTTCCCCGTCAAGCTCTAAATCGGGGGCTCCCTTTAGGGTTCCGATTTAGTGCTTTACGGCACCTCGACCCCAAAAAACTTGATTAGGGTGATG
GTTCACGTAGTGGGCCATCGCCCTGATAGACGGTTTTTCGCCCTTTGACGTTGGAGTCCACGTTCTTTAATAGTGGACTCTTGTTCCAAACTGGA
ACAACACTCAACCCTATCTCGGTCTATTCTTTTGATTTATAAGGGATTTTGCCGATTTCGGCCTATTGGTTAAAAAATGAGCTGATTTAACAAAA
ATTTAACGCGAATTTTAACAAAATATTAACGCTTACAATTTAGGTGGCACTTTTCGGGGAAATGTGCGCGGAACCCCTATTTGTTTATTTTTCTA
AATACATTCAAATATGTATCCGCTCATGAATTAATTCTTAGAAAAACTCATCGAGCATCAAATGAAACTGCAATTTATTCATATCAGGATTATCA
ATACCATATTTTTGAAAAAGCCGTTTCTGTAATGAAGGAGAAAACTCACCGAGGCAGTTCCATAGGATGGCAAGATCCTGGTATCGGTCTGCGAT
TCCGACTCGTCCAACATCAATACAACCTATTAATTTCCCCTCGTCAAAAATAAGGTTATCAAGTGAGAAATCACCATGAGTGACGACTGAATCCG
GTGAGAATGGCAAAAGTTTATGCATTTCTTTCCAGACTTGTTCAACAGGCCAGCCATTACGCTCGTCATCAAAATCACTCGCATCAACCAAACCG
TTATTCATTCGTGATTGCGCCTGAGCGAGACGAAATACGCGATCGCTGTTAAAAGGACAATTACAAACAGGAATCGAATGCAACCGGCGCAGGAA
CACTGCCAGCGCATCAACAATATTTTCACCTGAATCAGGATATTCTTCTAATACCTGGAATGCTGTTTTCCCGGGGATCGCAGTGGTGAGTAACC
ATGCATCATCAGGAGTACGGATAAAATGCTTGATGGTCGGAAGAGGCATAAATTCCGTCAGCCAGTTTAGTCTGACCATCTCATCTGTAACATCA
TTGGCAACGCTACCTTTGCCATGTTTCAGAAACAACTCTGGCGCATCGGGCTTCCCATACAATCGATAGATTGTCGCACCTGATTGCCCGACATT
ATCGCGAGCCCATTTATACCCATATAAATCAGCATCCATGTTGGAATTTAATCGCGGCCTAGAGCAAGACGTTTCCCGTTGAATATGGCTCATAA
CACCCCTTGTATTACTGTTTATGTAAGCAGACAGTTTTATTGTTCATGACCAAAATCCCTTAACGTGAGTTTTCGTTCCACTGAGCGTCAGACCC
CGTAGAAAAGATCAAAGGATCTTCTTGAGATCCTTTTTTTCTGCGCGTAATCTGCTGCTTGCAAACAAAAAAACCACCGCTACCAGCGGTGGTTT

| | |
|---|---|
| | GTTTGCCGGATCAAGAGCTACCAACTCTTTTTCCGAAGGTAACTGGCTTCAGCAGAGCGCAGATACCAAATACTGTCCTTCTAGTGTAGCCGTAG<br>TTAGGCCACCACTTCAAGAACTCTGTAGCACCGCCTACATACCTCGCTCTGCTAATCCTGTTACCAGTGGCTGCTGCCAGTGGCGATAAGTCGTG<br>TCTTACCGGGTTGGACTCAAGACGATAGTTACCGGATAAGGCGCAGCGGTCGGGCTGAACGGGGGGTTCGTGCACACAGCCCAGCTTGGAGCGAA<br>CGACCTACACCGAACTGAGATACCTACAGCGTGAGCTATGAGAAAGCGCCACGCTTCCCGAAGGGAGAAAGGCGGACAGGTATCCGGTAAGCGGC<br>AGGGTCGGAACAGGAGAGCGCACGAGGGAGCTTCCAGGGGGAAACGCCTGGTATCTTTATAGTCCTGTCGGGTTTCGCCACCTCTGACTTGAGCG<br>TCGATTTTTGTGATGCTCGTCAGGGGGGCGGAGCCTATGGAAAAACGCCAGCAACGCGGCCTTTTTACGGTTCCTGGCCTTTTGCTGGCCTTTTG<br>CTCACATGTTCTTTCCTGCGTTATCCCCTGATTCTGTGGATAACCGTATTACCGCCTTTGAGTGAGCTGATACCGCTCGCCGCAGCCGAACGACC<br>GAGCGCAGCGAGTCAGTGAGCGAGGAAGCGGAAGAGCGCCTGATGCGGTATTTTCTCCTTACGCATCTGTGCGGTATTTCACACCGCATATATGG<br>TGCACTCTCAGTACAATCTGCTCTGATGCCGCATAGTTAAGCCAGTATACACTCCGCTATCGCTACGTGACTGGGTCATGGCTGCGCCCCGACAC<br>CCGCCAACACCCGCTGACGCGCCCTGACGGGCTTGTCTGCTCCCGGCATCCGCTTACAGACAAGCTGTGACCGTCTCCGGGAGCTGCATGTGTCA<br>GAGGTTTTCACCGTCATCACCGAAACGCGCGAGGCAGCTGCGGTAAAGCTCATCAGCGTGGTCGTGAAGCGATTCACAGATGTCTGCCTGTTCAT<br>CCGCGTCCAGCTCGTTGAGTTTCTCCAGAAGCGTTAATGTCTGGCTTCTGATAAAGCGGGCCATGTTAAGGGCGGTTTTTTCCTGTTTGGTCACT<br>GATGCCTCCGTGTAAGGGGGATTTCTGTTCATGGGGGTAATGATACCGATGAAACGAGAGAGGATGCTCACGATACGGGTTACTGATGATGAACA<br>TGCCCGGTTACTGGAACGTTGTGAGGGTAAACAACTGGCGGTATGGATGCGGCGGGACCAGAGAAAAATCACTCAGGGTCAATGCCAGCGCTTCG<br>TTAATACAGATGTAGGTGTTCCACAGGGTAGCCAGCAGCATCCTGCGATGCAGATCCGGAACATAATGGTGCAGGGCGCTGACTTCCGCGTTTCC<br>AGACTTTACGAAACACGGAAACCGAAGACCATTCATGTTGTTGCTCAGGTCGCAGACGTTTTGCAGCAGCAGTCGCTTCACGTTCGCTCGCGTAT<br>CGGTGATTCATTCTGCTAACCAGTAAGGCAACCCCGCCAGCCTAGCCGGGTCCTCAACGACAGGAGCACGATCATGCGCACCCGTGGGGCCGCCA<br>TGCCGGCGCCTGCCACCATACCCACGCCGAAACAAGCGCTCATGAGCCCGAAGTGGCGAGCCCGATCTTCCCCATCGGTGATGTCGGCGATATAG<br>GCGCCAGCAACCGCACCTGTGGCGCCGGTGATGCCGGCCACGATGCGTCCGGCGTAGAGGATCGAGATCTCGATCCCGCGAAATTAATACGACTC<br>ACTATAGGGGAATTGTGAGCGGATAACAATTCCCTTCTAGCCATATTGGACTCGGACCTGTTTCACGTGGAACACTGAGCCTGGACTAGGTCTAG<br>CATATTGGACTCGACTTTCACGTGGAACACTGAGCCTGGACTAGGTCTAGAACAGGTTCTTTTTTCTTTGTTTCACGTGGAACATTCTGATTAAT<br>GTACAGCTAGCCATATTGGACTCGGACCTGTTTCACGTGGAACACTGAGCCTGGACTAGGTCTAGAACAGGTTCTTTTTTCTTTGTTTCACGTGG<br>AACATTCTGATTAATGTACAGCTAGAACAGGTTCTTTTTTCTTTGTTTCACGTGGAACATTCTGATTAATGTACAGCTAGAAATAATTTTGTTTA<br>ACTTTAAGAAGGAGATATACCATGTGTTTCACGTGGAACACATGGCTAAAGGCCTTGGAAAAGGGATTAATGCGTTATTTAATCAGGTAGATTTG<br>TCTGAAGAGACAGTTGAAGAAATTAAAATTGCCGATTTACGCCCTAATCCTTATCAGCCAAGAAAACACTTTGATGACGAGGCATTAGCTGAACT<br>AAAAGAATCTGTGCTGCAGCATGGCATTCTTCAGCCGCTTATCGTCAGAAAATCTTTAAAAGGCTATGATATTGTTGCGGGTGAACGGCGTTTTC<br>GAGCGGCAAAGCTGGCAGGTTTAGATACAGTTCCGGCCATTGTCCGTGAATTATCAGAGGCGTTAATGAGGGAAATTGCTTTATTAGAAAACCTT<br>CAGCGTGAAGATTTATCTCCGCTTGAAGAGGCTCAGGCATATGCCGAAACAAGCGCTCATGAGCCCGAAGTGGCGAGCCCGATCTTCCCCATCGG<br>TGATGTCGGCGATATAGGCGCCAGCAACCGCACCTGTGGCGCCGGTGATGCCGGCCACGATGCGTCCGGCGTAGAGGATCGAGATCTCGATCCCG<br>CGAAATTAATACGACTCACTATAGGGGAATTGTGAGCGGATAACAATTCCCTTCTAGCCATATTGGACTCGGACCTGTTTCACGTGGAACACTGA<br>GCCTGGACTAGGTCTAGCATATTGGACTCGACTTTCACGTGGAACACTGAGCCTGGACTAGGTCTAGAACAGGTTCTTTTTTCTTTGTTTCACGT<br>GGAACATTCTGATTAATGTACAGCTAGCCATATTGGACTCGGACCTGTTTCACGTGGAACACTGAGCCTGGACTAGGTCTAGAACAGGTTCTTTT<br>TTCTTTGTTTCACGTGGAACATTCTGATTAATGTACAGCTAGAACAGGTTCTTTTTTCTTTGTTTCACGTGGAACATTCTGATTAATGTACAGCT<br>AGAAATAATTTTGTTTAACTTTAAGAAGGAGATATACCATGTGTTTCACGTGGAACACATGGCTAAAGGCCTTGGAAAAGGGATTAATGCGCATA<br>TGACTCCCTTTTGAAACACTTAGATCTCACACAAGAGCAGCTTGCCAAACGTCTTGGGAAAAGCAGACCGCATATTGCGAATCATTTAAGACTGC<br>TGACACTGCCAGAAAATATTCAACAGCTTATTGCCGAAGGCACGCTTTCTATGGGACATGGACGCACGCTTCTTGGCTTAAAAAACAAAAATAAG<br>CTTGAACCGCTGGTACAAAAAGTGATTGCGGAGCAGCTCAATGTTCGCCAACTTGAGCAGCTGATTCAGCAGTTGAATCAGAATGTTCCACGTGA<br>AACAAAGAAAAAAGAACCTGTGAAAGATGCGGTTCGCCATGCCGGCGCCTGCCACCATACCCACGCCGAAACAAGCGCTCATGAGCCCGAAGTGG<br>CGAGCCCGATCTTCCCCATCGGTGATGTCGGCGATATAGGCGCCAGCAACCGCACCTGTGGCGCCGGTGATGCCGGCCACGATGCGTCCGGCGTA<br>GAGGATCGAGATCTCGATCCCGCGAAATTAATACGACTCACTATAGGGGAATTGTGAGCGGATAACAATTCCCTTCTAGCCATATTGGACTCGGA<br>CCTGTTTCACGTGGAACACTGAGCCTGGACTAGGTCTAGCATATTGGACTCGACTTTCACGTGGAACACTGAGCCTGGACTAGGTCTAGAACAGG<br>TTCTTTTTTCTTTGTTTCACGTGGAACATTCTGATTAATGTACAGCTAGCCATATTGGACTCGGACCTGTTTCACGTGGAACACTGAGCCTGGAC<br>TAGGTCTAGAACAGGTTCTTTTTTCTTTGTTTCACGTGGAACATTCTGATTAATGTACAGCTAGAACAGGTTCTTTTTTCTTTGTTTCACGTGGA<br>ACATTCTGATTAATGTACAGCTAGAAATAATTTTGTTTAACTTTAAGAAGGAGATATACCATGTGTTTCACGTGGAACACATGGCTAAAGGCCTT<br>GGAAAAGGGATTAATGCGTTATTTAATCAGGTAGATTTGTCTGAAGAGACAGTTGAAGAAATTAAAATTGCCGATTTACGCCCTAATCCTTATCA<br>GCCAAGAAAACACTTTGATGACGAGGCATTAGCTGAACTAAAAGAATCTGTGCTGCAGCATGGCATTCTTCAGCCGCTTATCGTCAGAAAATCTT<br>TAAAAGGCTATGATATTGTTGCGGGTGAACGGCGTTTTCGAGCGGCAAAGCTGGCAGGTTTAGATACAGTTCCGGCCATTGTCCGTGAATTATCA<br>GAGGCGTTAATGAGGGAAATTGCTTTATTAGAAAACCTTCAGCGTGAAGATTTATCTCCGCTTGAAGAGGCTCAGGCATATGCCGAAACAAGCGC<br>TCATGAGCCCGAAGTGGCGAGCCCGATCTTCCCCATCGGTGATGTCGGCGATATAGGCGCCAGCAACCGCACCTGTGGCGCCGGTGATGCCGGCC<br>ACGATGCGTCCGGCGTAGAGGATCGAGATCTCGATCCCGCGAAATTAATACGACTCACTATAGGGGAATTGTGAGCGGATAACAATTCCCTTCTA<br>GCCATATTGGACTCGGACCTGTTTCACGTGGAACACTGAGCCTGGACTAGGTCTAGCATATTGGACTCGACTTTCACGTGGAACACTGAGCCTGG<br>ACTAGGTCTAGAACAGGTTCTTTTTTCTTTGTTTCACGTGGAACATTCTGATTAATGTACAGCTAGCCATATTGGACTCGGACCTGTTTCACGTG<br>GAACACTGAGCCTGGACTAGGTCTAGAACAGGTTCTTTTTTCTTTGTTTCACGTGGAACATTCTGATTAATGTACAGCTAGAACAGGTTCTTTTT<br>TCTTTGTTTCACGTGGAACATTCTGATTAATGTACAGCTAGAAATAATTTTGTTTAACTTTAAGAAGGAGATATACCATGTGTTTCACGTGGAAC<br>ACATGGCTAAAGGCCTTGGAAAAGGGATTAATGCGCATATGACTCCCTTTTGAAACACTTAGATCTCACACAAGAGCAGCTTGCCAAACGTCTTG<br>GGAAAAGCAGACCGCATATTGCGAATCATTTAAGACTGCTGACACTGCCAGAAAATATTCAACAGCTTATTGCCGAAGGCACGCTTTCTATGGGA<br>CATGGACGCACGCTTCTTGGCTTAAAAAACAAAAATAAGCTTGAACCGCTGGTACAAAAAGTGATTGCGGAGCAGCTCAATGTTCGCCAACTTGA<br>GCAGCTGATTCAGCAGTTGAATCAGAATGTTCCACGTGAAACAAAGAAAAAAGAACCTGTGAAAGATGCGGTTCGCCATGCCGGCGATAATGGCC<br>TGCTTCTCGCCGAAACGTTTGGTGGCGGGACCAGTGACGAAGGCTTGAGCGAGGGCGTGCAAGATTCCGAATACCGCAAGCGACAGGCCGATCAT<br>CGTCGCGCTCCAGCGAAAGCGGTCCTCGCCGAAAATGACCCAGAGCGCTGCCGGCACCTGTCCTACGAGTTGCATGATAAAGAAGACAGTCATAA<br>GTGCGGCGACGATAGTCATGCCCCGCGCCCACCGGAAGGAGCTGACTGGGTTGAAGGCTCTCAAGGGCATCGGTCGAGATCCCGGTGCCTAATGA<br>GTGAGCTAACTTACATTAATTGCGTTGCGCTCACTGCCCGCTTTCCAGTCGGGAAACCTGTCGTGCCAGCTGCATTAATGAATCGGCCAACGCGC<br>GGGGAGAGGCGGTTTGCGTATTGGGCGCCAGGGTGGTTTTTCTTTTCACCAGTGAGACGGGCAACAGCTGATTGCCCTTCACCGCCTGGCCCTGA<br>GAGAGTTGCAGCAAGCGGTCCACGCTGGTTTGCCCCAGCAGGCGAAAATCCTGTTTGATGGTGGTTAACGGCGGGATATAACATGAGCTGTCTTC<br>GGTATCGTCGTATCCCACTACCGAGATATCCGCACCAACGCGCAGCCCGGACTCGGTAATGG |

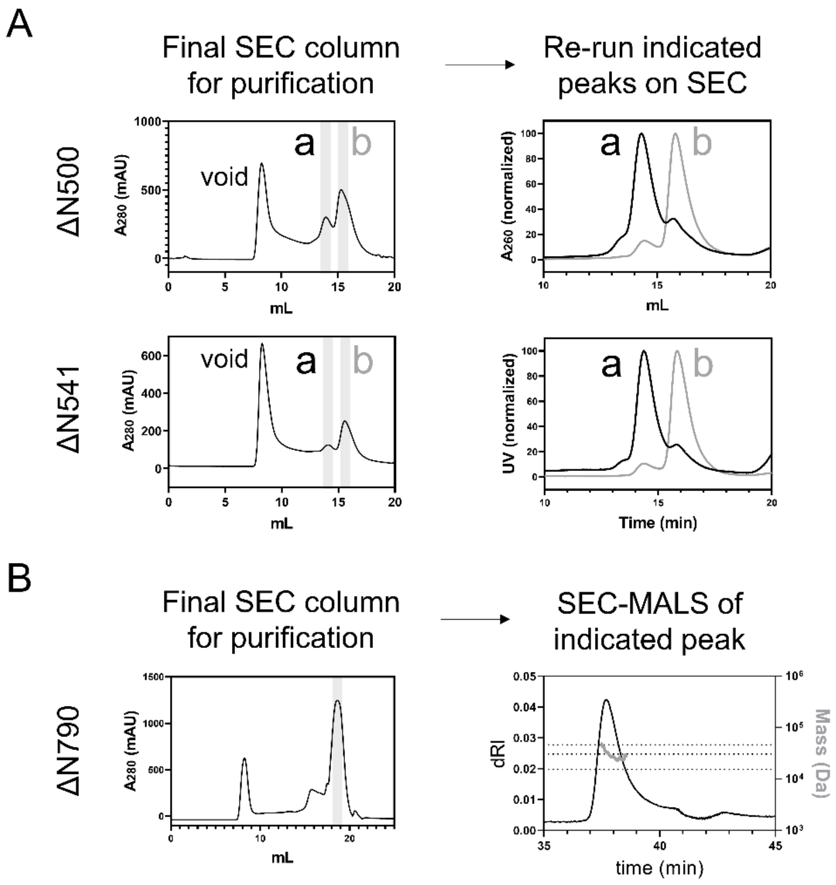


**Supplementary Figure 1.** ***SEC and SEC-MALS analyses of ΔN variants.*** (A) The ΔN500 and ΔN541 variants both ran as two peaks (a and b) during the final size exclusion chromatography (SEC) run during purification. When selected peaks were re-run on SEC columns, they did not fully re-equilibrate into the two original peaks. In combination with MALS analysis (see main text) this suggests that these variants exist as a slowly exchanging population of monomers and dimers. (B) The ΔN4 variant was purified by SEC and then subjected to SEC-MALS analysis. The data quality was poor, yielding equivocal molecular weights ranging from 1.5 to 3.1x the value expected for a ΔN4 monomer. The dotted lines on the graph indicated the expected masses for a 1mer, 2mer and 3mer. Therefore, this variant was also analysed by native mass spectrometry which showed it was monomeric (see main text).

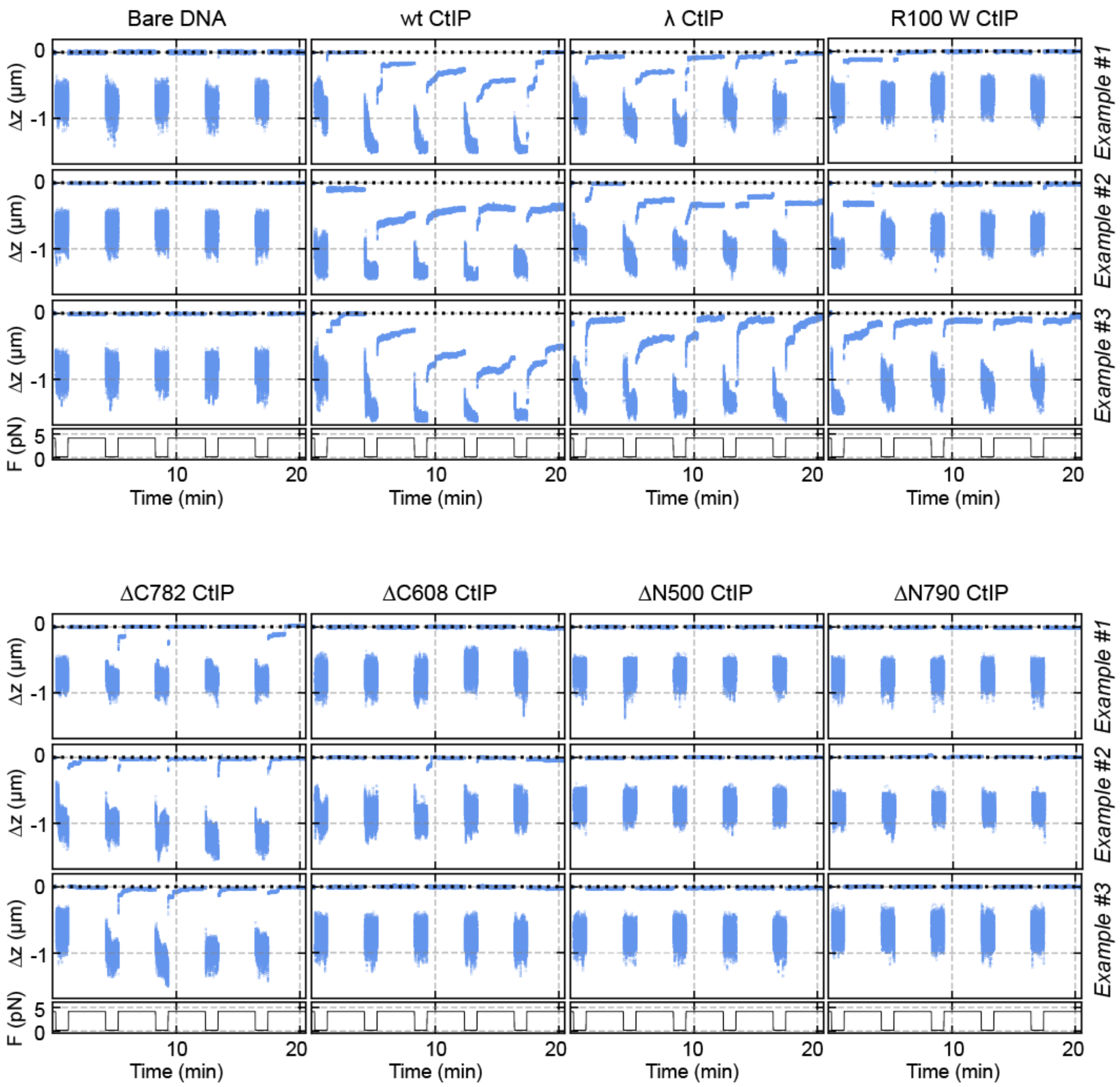


**Supplementary Figure 2.** ***Representative time-courses from the magnetic tweezers' experiments.*** Individual time courses for the different wild-type and mutant CtIP variations studied in this work. See methods for a detailed description of the magnetic tweezers force-cycle assay.

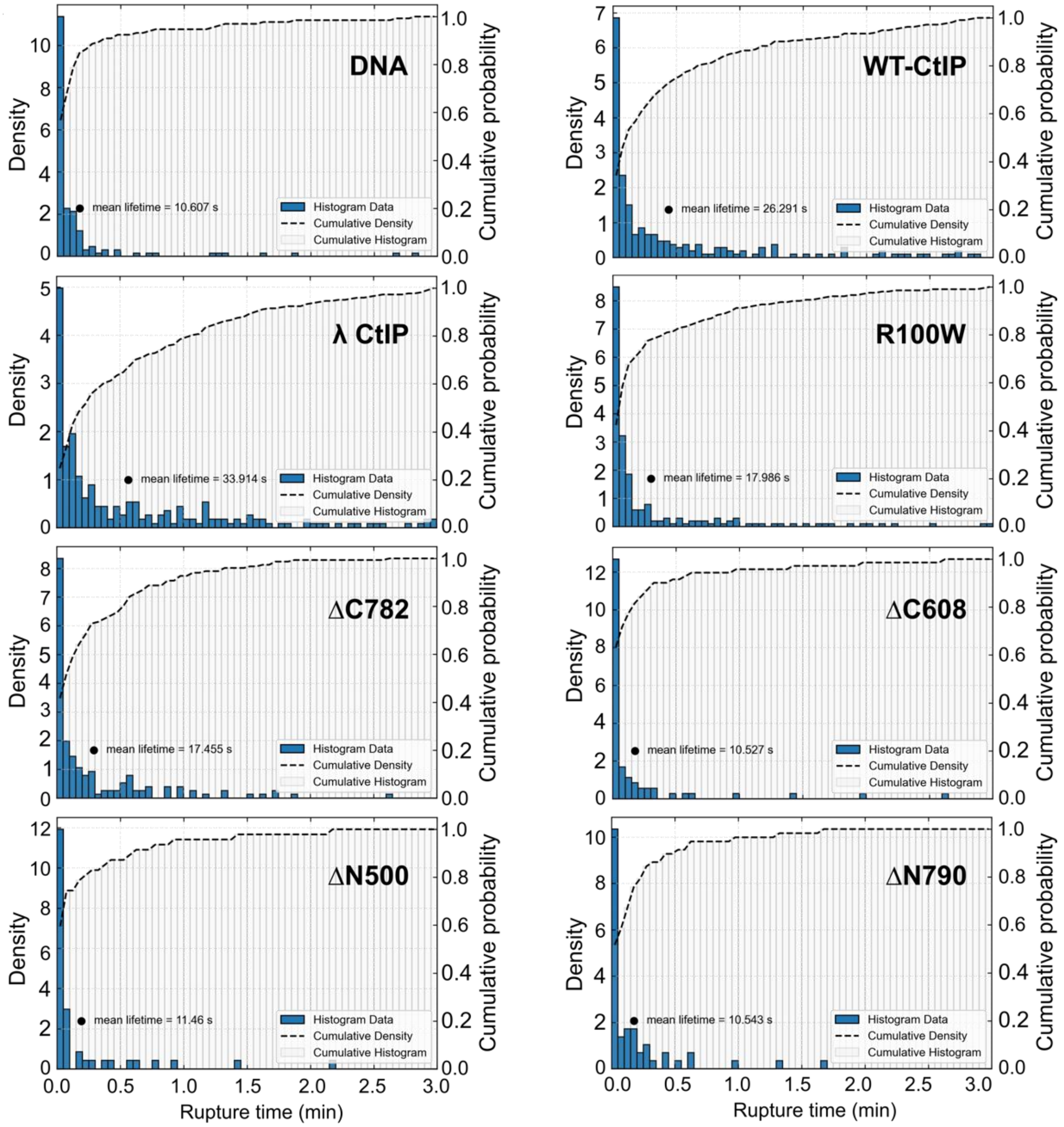


**Supplementary Figure 3.** ***Lifetime distributions for Class II events.*** Each panel represents the distribution of the rupture time, the cumulative probability of rupture and the average lifetime before rupture for each CtIP variant studied.

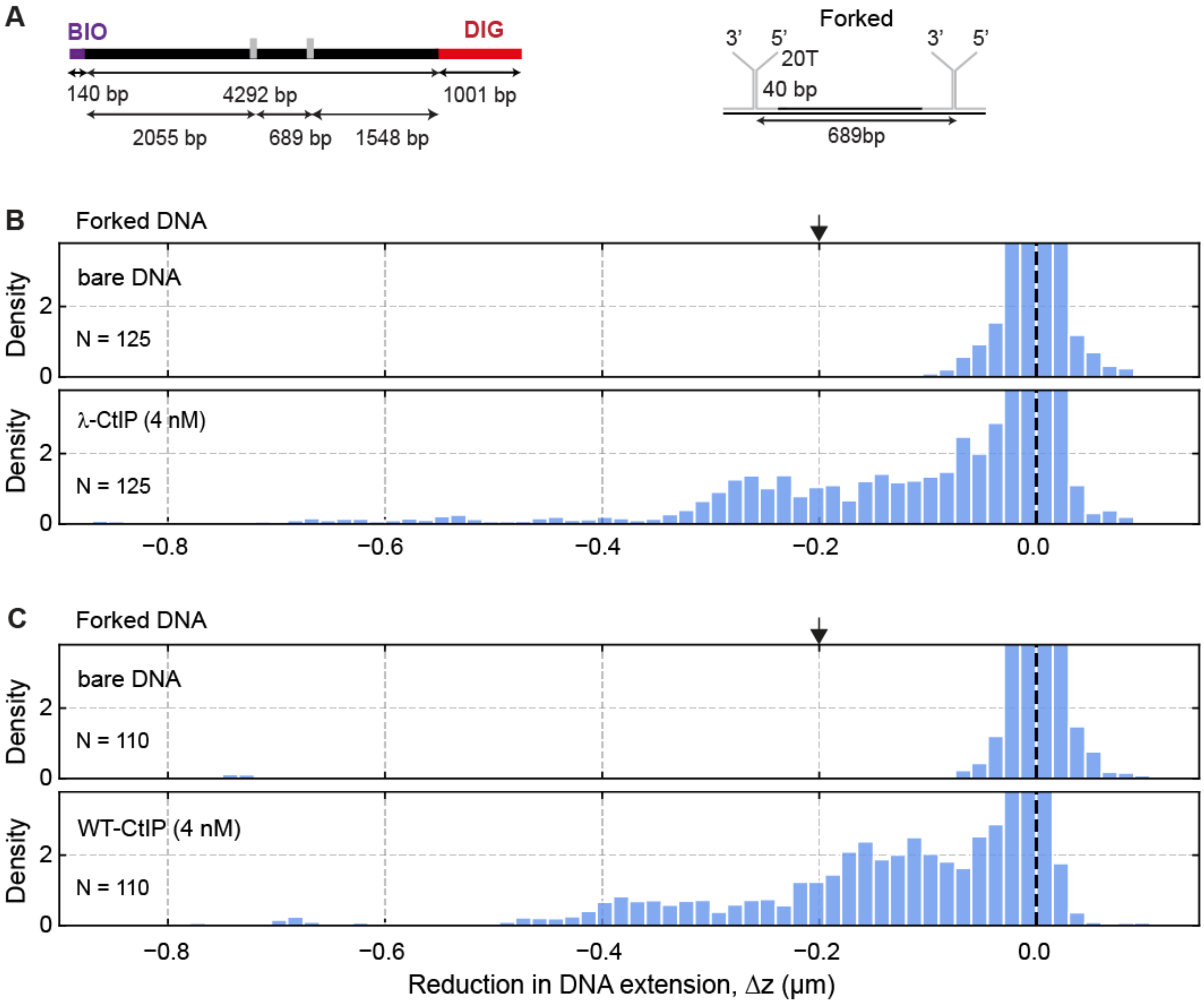


**Supplementary Figure 4.** ***Bridging interactions do not specifically tether DNA ends.*** (**A**) Cartoon of a branched DNA substrate with forked ends to study dsDNA bridging with MT. The right panel is a zoom of the branched region. Not to scale. (**B**) Histogram of the relative reductions (Δz) for DNA molecules containing two forked branches in the absence of protein (upper panel) and in the presence of 4 nM λ-CtIP (lower panel). (**C**) Histogram of the relative reductions (Δz) for DNA molecules containing two forked branches in the absence of protein (upper panel) and in the presence of 4nM WT-CtIP (lower panel). The peak at $\Delta z = 0$ µm represents all extended states and the densities for $\Delta z < 0$ µm represent all bridged states through CtIP interactions. Arrows indicate the expected reduction for a specific end-to-end bridging between both fork ends. N indicates the number of DNA molecules studied for each condition.

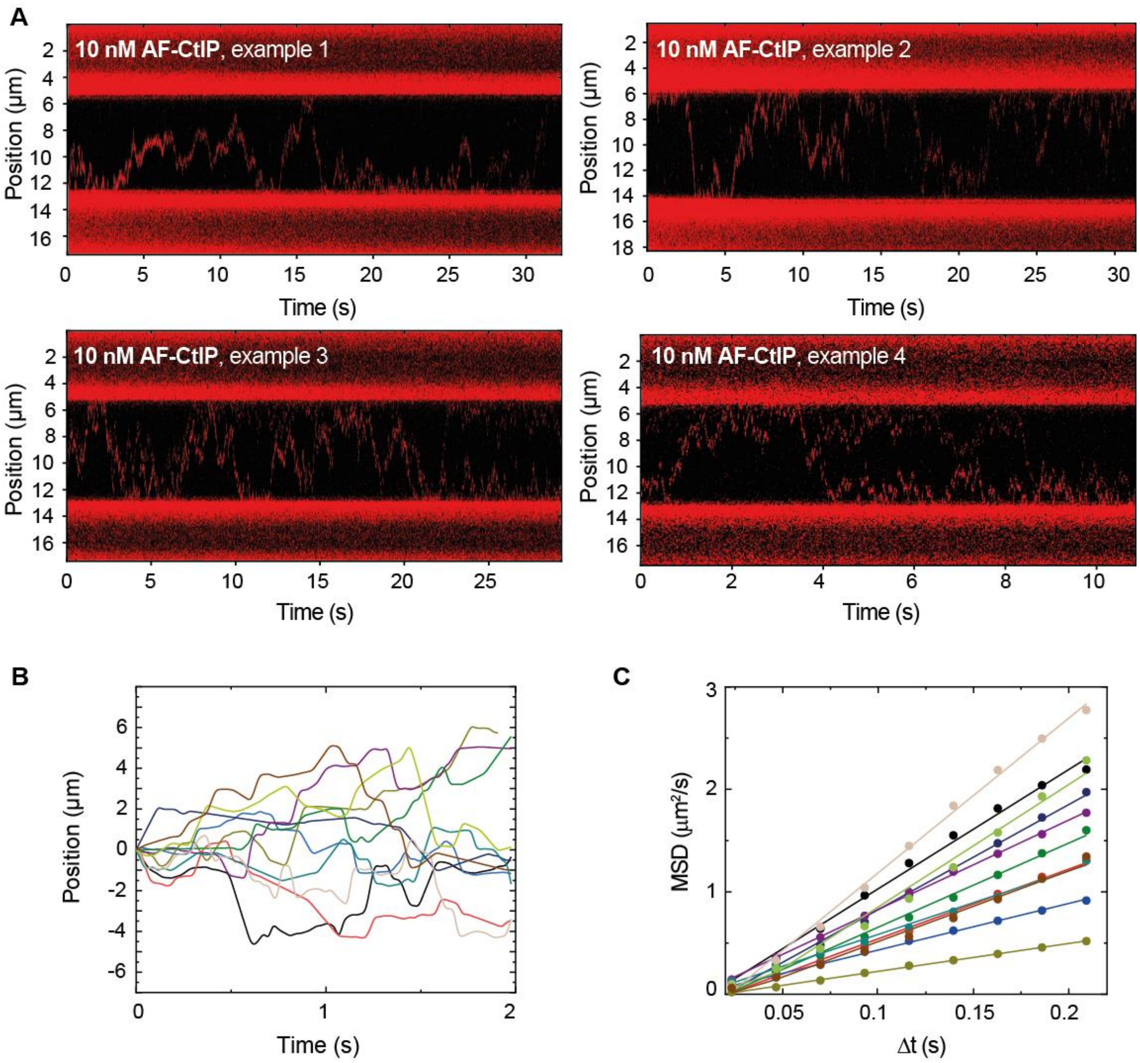


**Supplementary Figure 5.** ***Representative kymographs of individual trajectories of CtIP diffusion on DNA.*** **(A)** Representative examples of kymographs showing trajectories of individual WT-CtIP proteins labelled with Alexa Fluor 635 (AF). (**B**) Position of CtIP over time determined from the analysis of CtIP kymographs (N =115). (**C**) Mean squared displacement (MSD) of CtIP for different time intervals (Δt). Straight lines indicate normal (i.e., random walk) diffusive behaviour.
